# The natural history of bacterial bloomers in a decade-long time series

**DOI:** 10.64898/2026.07.01.735832

**Authors:** Ona Deulofeu-Capo, Carmen García-Comas, Xavier Rey-Velasco, Adrià Auladell, Ramiro Logares, Esther Garcés, Isabel Ferrera, Olga Sánchez, Josep M Gasol, Marta Sebastián

## Abstract

Bacterial bloomers—populations that experience rapid and significant increases in abundance in response to environmental triggers—briefly dominate marine microbial communities, potentially impacting the ecosystem by channeling large amounts of nutrients and affecting carbon fluxes. Due to their ephemeral nature, bacterial bloomers are challenging to capture, and it remains unknown whether they are restricted to specific taxonomic groups or whether they exhibit recurrent patterns. We analyzed a decade-long time series from the Blanes Bay Microbial Observatory (BBMO, NW Mediterranean Sea) to investigate bacterial bloomers in two size fractions (free-living (0.2–3 µm) and particle-attached (3–20 µm) communities. We identified 57 Amplicon Sequence Variants (ASVs), less than 1% of the total bacterial richness, exhibiting recurrent or chaotic blooming-like behavior. Bloomers spanned diverse phyla, though some taxonomic coherence appeared within families containing multiple blooming taxa. Monthly sampling detected bloom events on average 4.6 ± 1.9 times per year across both size fractions. Once seasonality was accounted for, blooms showed weak associations with biological and physicochemical variables, likely a consequence of monthly sampling resolution. Nonetheless, a marked shift in the blooming community within the particle-attached size fraction coincided with ecosystem disturbances from the nearby harbour restoration, suggesting that bloomers may act as disturbance sentinels. Metagenomic data showed that blooms led to marked shifts in the community functional potential. Overall, our findings underscore the importance of investigating bloom dynamics to understand microbial contributions to biogeochemical cycles and stress the need for higher-frequency sampling to accurately capture these transient but ecologically relevant events.

## Introduction

Marine microbial communities comprise thousands of different taxa that undergo continuous succession in response to environmental fluctuations [1], with substantial variations across both space [2] and time [3]. Occasionally, certain taxa, the so-called “bloomers”, experience a sharp increase in abundance, triggering a blooming event that leads to abrupt shifts in community structure [4–8], potentially channeling a substantial fraction of carbon or nutrient fluxes [9]. Blooming events result from the complex interplay of abiotic factors (e.g., nutrient availability, temperature), biotic interactions (e.g., competition, grazing), and intrinsic traits that enable certain taxa to bloom, collectively shaping community dynamics [10]. However, although blooms of phototrophic eukaryotes and prokaryotes, such as diatoms or cyanobacteria, are well studied with well-documented impacts on community function [11–17], blooms of heterotrophic prokaryotes and their ecological consequences remain largely unexplored.

Previous research on blooming heterotrophic bacteria has primarily focused on their association with phytoplankton blooms [12, 17–21]. However, short-lived increases in certain fast-growing taxa (i.e. blooms) have occasionally been reported in aquatic ecosystems, without clear links to environmental factors [22–24]. These observations align with the view that nearly all environments harbor taxa capable of abrupt abundance shifts, often playing a disproportionately significant role in shaping overall community dynamics [7]. Despite their potential ecological significance, these events have historically been overlooked, likely due to the challenge of capturing their ephemeral nature with regular sampling programs and because detecting bacterial blooms requires molecular approaches.

Long-term microbial observatories offer valuable insights into the structure and function of microbial communities. Although the sampling frequency of these observatories is typically low (e.g., monthly), their extended duration increases the likelihood of capturing blooming events, to study their frequency, recurrence and their potential impact; knowledge essential for improving our predictions of community shifts impact on biogeochemical cycles. Understanding blooming dynamics requires addressing fundamental questions, including how to define a bacterial bloomer, to determine the frequency of blooms, assess the role of environmental conditions and community interactions in shaping bloom events, and explore their potential impact. To address these questions, we used a 10-year monthly metabarcoding time series in the NW Mediterranean Sea. Our specific objectives were: (1) to identify bloomers in the time series, (2) to determine whether they exhibit recurrence, (3) to assess whether bloom-like behavior is taxonomically coherent, (4) to explore the environmental and biological variables causally affecting them, and (5) to examine if blooming events are linked to functional shifts in the microbial community.

## Materials and methods

Surface water pre-sieved through a 200-μm mesh was collected in 20-L plastic carboys from the Blanes Bay Microbial Observatory (BBMO), a shallow (∼20 m) coastal station ∼1 km offshore in the Northwestern Mediterranean (41°40’N, 2°48’E), and transported to the lab within 2 hours. The dataset studied corresponds to ten years (2004-2013) of monthly sampling whose seasonal changes in environmental parameters have been extensively characterized [25, 26]. The following variables were measured: water temperature (°C) and salinity, day-length (hours of light), transparency (Secchi depth, in m), the concentration of inorganic nutrients, abundances of heterotrophic bacteria, photosynthetic picophytoplankton, heterotrophic and phototrophic nanoflagellates, total and <3 µm chlorophyll-*a* concentration and bacterial heterotrophic activity (see Supplementary Information I for details on the methods). The previously mentioned variables, when missing, were interpolated using the LOESS function (*stats* package, base R) only if the gaps were non-consecutive and accounted for less than 10% of the time series. We calculated a synthetic Euclidean distance from consecutive samplings, with variables normalized to z-scores, to summarize changes in environmental conditions separately for physicochemical and biological variables.

### DNA extraction and sequencing

Samples were sieved through a 20-μm mesh to remove large particles and about 4-L of microbial biomass was concentrated, using a peristaltic pump, onto a 3-μm polycarbonate filter for the 3-20 μm size fraction, including particle-attached bacteria, and then filtered through a 0.2-μm Sterivex unit (Millipore) for the 0.2-3 μm size fraction, including free-living bacteria. Details on DNA extraction and sequencing can be found in Supplementary Information I.

### Identification of bloomers and definition of blooming events

To identify bloomers in the time series, we considered three criteria: (1) the bloomer’s relative abundance had to exceed 10% of the total community at blooming time point; (2) at that identified time point, the z-score (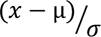) of its abundance (x) considering the mean (µ) between that point and the preceding two timepoints divided by the standard deviation (*σ*), had to be greater than 2 (when relative abundance was considered) and 1 (when robust centered log ratio (rCLR) was considered), and (3) the coefficient of variation (cv) of the relative abundance values throughout the time series had to be greater than 1 (i.e. indicating dynamic behavior). We used both, relative abundance and rCLR transformed abundances to overcome artifacts in the bloomers detection due to the compositional nature of the data [27]. Since bloomer detection was robust to z-score variation (1, 2 and 10; Fig. S1), we applied z-score thresholds of 1 and 2 based on relative abundance and rCLR. The relative abundance threshold, conversely, was the primary driver of bloomer detection sensitivity. We set this threshold at 10% as such abundant ASV are more likely to impact community function.

### Classification of bloomers

Bloomers were classified based on their occurrence (i.e. number of time points in which an ASV was detected) and their recurrence (i.e., patterns of abundance periodicity). Based on occurrence, three categories were defined: narrow (occurrence in < 1/3 of the samples), intermediate (1/3 - 2/3 occurrence) and broad (> 2/3 occurrence). Recurrence was assessed using wavelet analysis on each bloomer-ASV time series. Wavelet transformation is well suited for detecting periodicity and changes in frequency over time, providing a multi-scale representation of temporal patterns and enabling identification of recurrent abundance dynamics [28, 29] (see Fig. S2 for examples of this analysis). Before analysis, we verified equidistant sampling and transformed reads to rCLR. The three missing samples in the 3-20 μm size fraction, were linearly interpolated from adjacent measurements and visually checked for potential artifacts. Wavelets analysis was carried out using the package *WaveletComp* [30]. Non-recurrent bloomers were those exhibiting irregular (non-periodic) oscillations and have sometimes been referred to as ‘non-rhythmic’ [31]. Here we decided to term them ‘chaotic’ to distinguish from seasonal bloomers, as their irregular bloom dynamics are contingent on ecological initial conditions rather than periodic environmental cycles [32].

### Diversity analysis

To compute diversity, we used matrix normalization through rarefaction to the minimum number of reads per sample (N= 6,135) and repeated the step 1,000 times to overcome the uneven sampling effort bias with rrarefy.perm from *EcolUtils* package [33]. We calculated Bray-Curtis dissimilarity between pairs of contiguous samples as a measure for ß-diversity indicating temporal species turnover. We also calculated community evenness, as a measure of equality of abundances of ASVs in a community. We then checked if ß-diversity and evenness were affected during blooming events.

### Phylogenetic tree and phylogenetic distance

To assess whether bloom-like behavior is a taxonomically coherent trait, we constructed a phylogenetic tree with all ASVs in our dataset using raxml-ng/0.9.0 from an alignment produced by clustal/1.2.4. To analyze the bloomers relationships with their closely related taxa, we compared their sequences using the p-distance, i.e. the proportion of nucleotide sites at which the two sequences compared are different. We considered two ASVs as phylogenetically closely related when the p-distance was < 0.012, equivalent to 5 nucleotide differences between the ASV sequences [14, 34].

### Empirical dynamical model (EDM): convergent cross mapping (CCM) and Multiview distance regularized (MDR) S-map

To explore the environmental and biological variables causally influencing the broadly occurrent bloomers, we applied empirical dynamical modelling (EDM) [35, 36]. This is a powerful method that can help distinguish causality from spurious pair-wise correlations in time series from dynamical systems. EDM is a nonparametric method that reconstructs the dynamics of a system by embedding time-lagged time series of its components (i.e., state-space reconstruction from ASVs abundances). Thus, the method does not assume any equation governing the system, which makes it ideal to study non-linear context-dependent dynamics in microbial communities.

To reconstruct the state space from time-lagged embedding, EDM performs reliably only on those ASVs present in at least 2/3 of the samples, which included 5 and 7 bloomers in the 0.2-3 μm and 3-20 μm size fractions, respectively. Of the various tools within EDM, we applied Convergent Cross Mapping analysis (CCM) to test causality between pairs of variables [35, 36]. CCM tests if the reconstructed state space of an effect variable can recover (predict) the states of a potential causal variable. We applied CCM to pairs of variables including rCLR-transformed ASVs abundances and environmental variables. Prior to CCM analysis, we transformed the data to their monthly z-scores to remove the strong seasonal effect in Blanes and control for the Moran effect (i.e., synchronization of populations due to common external forcing). Once we knew ‘who affected who’, we applied Multiview Distance Regularised (MDR) S-map [37] on the rCLR-transformed abundances to infer the sign and strength of the interactions exerted and received by an ASVs from the community through time. This inference was possible through multiple state-space reconstructions for each ASV using time lags (up to three) from a subsample of causally related ASVs. The CCM was computed with the package *rEDM* [38] and the MDR S-map with: https://github.com/biozoo/MDR_S-map.

### Metagenomic data analysis

To examine if blooming events are linked to functional shifts in the microbial community and sustain relevant functions during blooms, we used a metagenomic dataset of the 0.2–3 μm size fraction collected between 2009 and 2013 [39]. Metagenome data processing is explained in Supplementary Information I. To explore whether changes in community composition were also evident at the genetic and functional level we calculated the Bray-Curtis dissimilarity between consecutive sampling points in the genetic composition (Single copy gene (SCG)-normalized) and the functional composition (i.e. KOs groups composition, SCG-normalized), and then assessed their correlation with the Bray-Curtis dissimilarity between consecutive sampling points of the metabarcoding community composition. The SCG-normalization provides a common scale across samples and serves as proxy for gene copy number per cell, enabling calculation of genetic and functional Bray–Curtis dissimilarities and bloomers’ specific contributions to KEGG functions. Since KOs group ORFs with similar functions, this approach provides a more ecologically meaningful perspective. Due to computational limitations, the gene-based dataset to calculate Bray-Curtis dissimilarity was reduced to include only genes with at least one abundance value exceeding 0.0003 over all the time series.

To explore possible functional changes linked to the different bloomers, we examined shifts in the total abundance (TPM-normalized) of genes sharing the same taxonomy with the identified bloomers. We analyzed functional shifts of three heterotrophic bloomers: *Glaciecola*–*ASV11*, *Amylibacter*–ASV27 and NS4 marine group –ASV58, whose taxonomically assigned genes in GTBD taxonomy belonged to: *Glaciecola*, *Amylibacter* and MAG121220-bin8, respectively. We assessed whether these blooming taxa contributed disproportionately to specific functions (KOs) during bloom events using the KOs-SCG normalized tables and calculated the relative contribution of each bloomer to each KO (i.e., the proportion of genes from a given bloomer assigned to a particular KO relative to the total KO count in the community).

### Data visualization and statistical analyses

All analyses were run in R v.4.2.3 [40] and Rstudio v.2022.12.0+353 [41] using *tidyverse* v.2.0.0 [42]. Plots were created with *ggplot2* v.3.4.4 [43] and *cowplot* v.1.1.1 [44]. Statistical differences between groups were assessed using the Kruskal-Wallis test due to non-normality (Shapiro-Wilk test, p < 0.01) and lack of homoscedasticity (Levene’s test, p < 0.01). Post-hoc comparisons were conducted using Dunn’s test to identify significant differences between groups (p < 0.01).

## Results

The BBMO microbial community exhibited interannual recurrence (Fig. 1A), with moderate total bacterial abundances variation over time (8.33·10^5^± 2.74·10^5^ (mean±SD; N= 120)) (Fig. S3A, Table S1), and a characteristic annual spring phytoplankton bloom (Fig. S3B). Despite this overall abundance stability, community structure varied markedly on a monthly scale through the time series, as reflected by the Bray-Curtis dissimilarity (Fig. 1B). Notably, some abrupt shifts in community composition did not co-occur with major environmental changes (Fig. 1C; Pearson correlation p-values>0.1).

**Figure 1.**
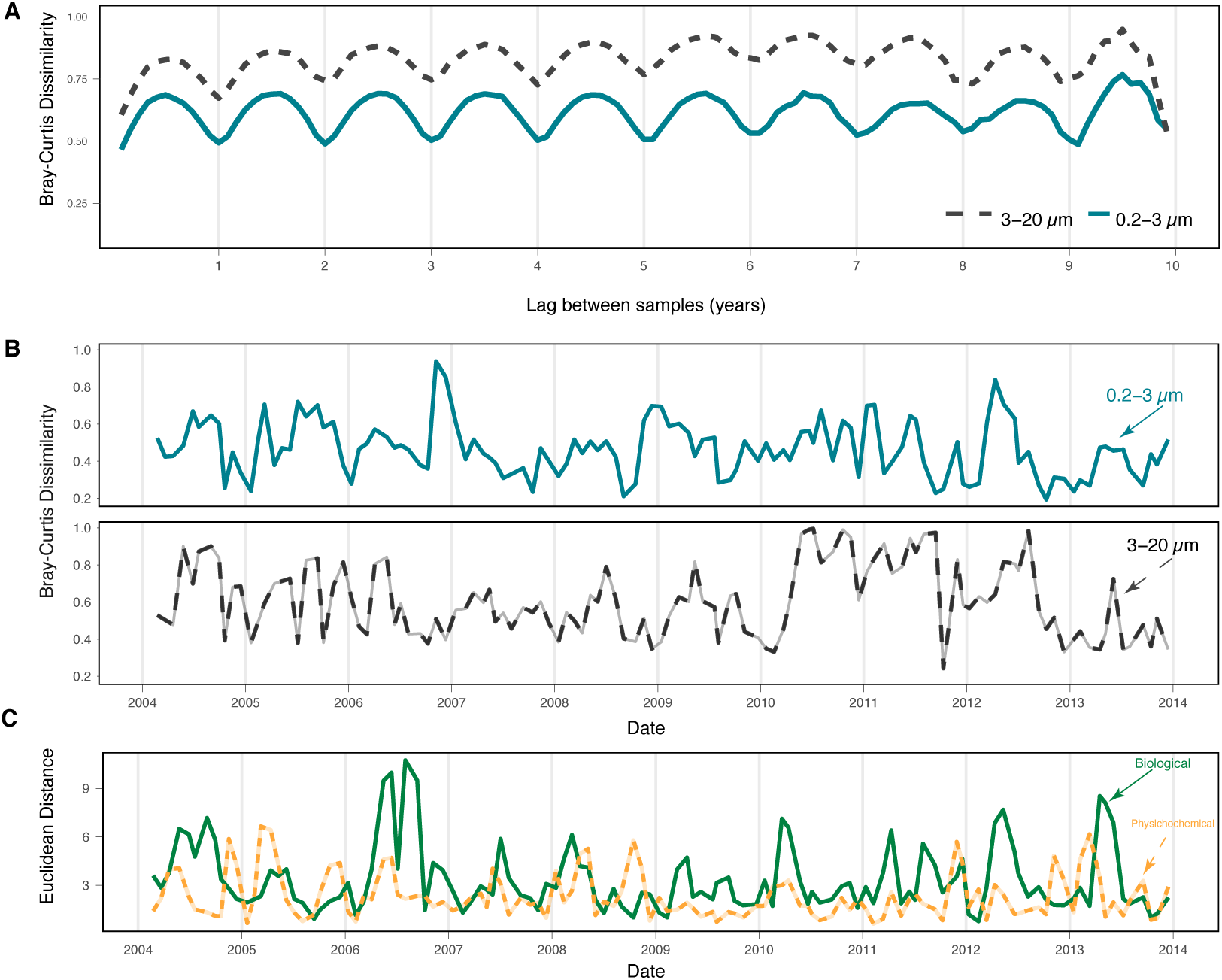
Summary of the 10-year time series from the Blanes Bay Microbial Observatory (BBMO): A) Recurrence over the years, measured by Bray-Curtis dissimilarity (Y-axis) across different time lags (X-axis). Solid line represents the average diversity values for the 0.2-3 µm (free-living) size fraction (circles), while the dashed line indicates the values for the 3-20 µm (particle-attached) size fraction (triangles). B) ß-diversity values over time for both size fractions: Bray-Curtis dissimilarity of community composition between pairs of consecutive time points. C) Synthetic Euclidean environmental distances between pairs of consecutive time points, calculated on physicochemical (inorganic nutrients (phosphate, silicate, nitrate, nitrite, and ammonia), temperature, and day length) (solid line) and biological (flow cytometry counts (prokaryotes and cyanobacteria), chlorophyll-*a* concentration, and the concentrations of heterotrophic and phototrophic nanoflagellates, *Cryptomonas*, and *Micromonas-like flagellates*) (dashed line) variables.

### Bacterial blooms in the 10y time series and recurrence patterns

First, based on our bloomer definition, we identified a total of 57 apparent bacterial bloomers in the 10y BBMO time series: 14 in the 0.2-3 μm fraction, 41 in the 3-20 μm fraction and 6 in both size fractions. Although they are only apparent bloomers due to the monthly sampling frequency of the time series, they will hereafter be referred to as bloomers. While bloomers accounted for less than 1% of the total community richness (n = 7849), their contribution to relative abundance was occasionally substantial. In some cases, blooming taxa represented up to 75% of the total bacterial abundance. On average, they accounted for 24.0% ± 14.2 of the 0.2–3 μm fraction and 37.1% ± 18.9 of the 3–20 μm fraction (Fig. 2A). Bloom events led to shifts in community composition, reflected by higher β-diversity values between consecutive samples (Fig. 2C), and a general decrease in community evenness (Fig. 2D). At the monthly scale sampling resolution, blooming events were detected with a mean frequency of 3.7±1.34 y^-1^ in the 0.2-3 μm fraction and 5.6±1.96 y^-1^ in the 3-20 μm fraction (Table S2, see Fig. S4 and Fig. S5 to visualize the bloom events).

**Figure 2.**
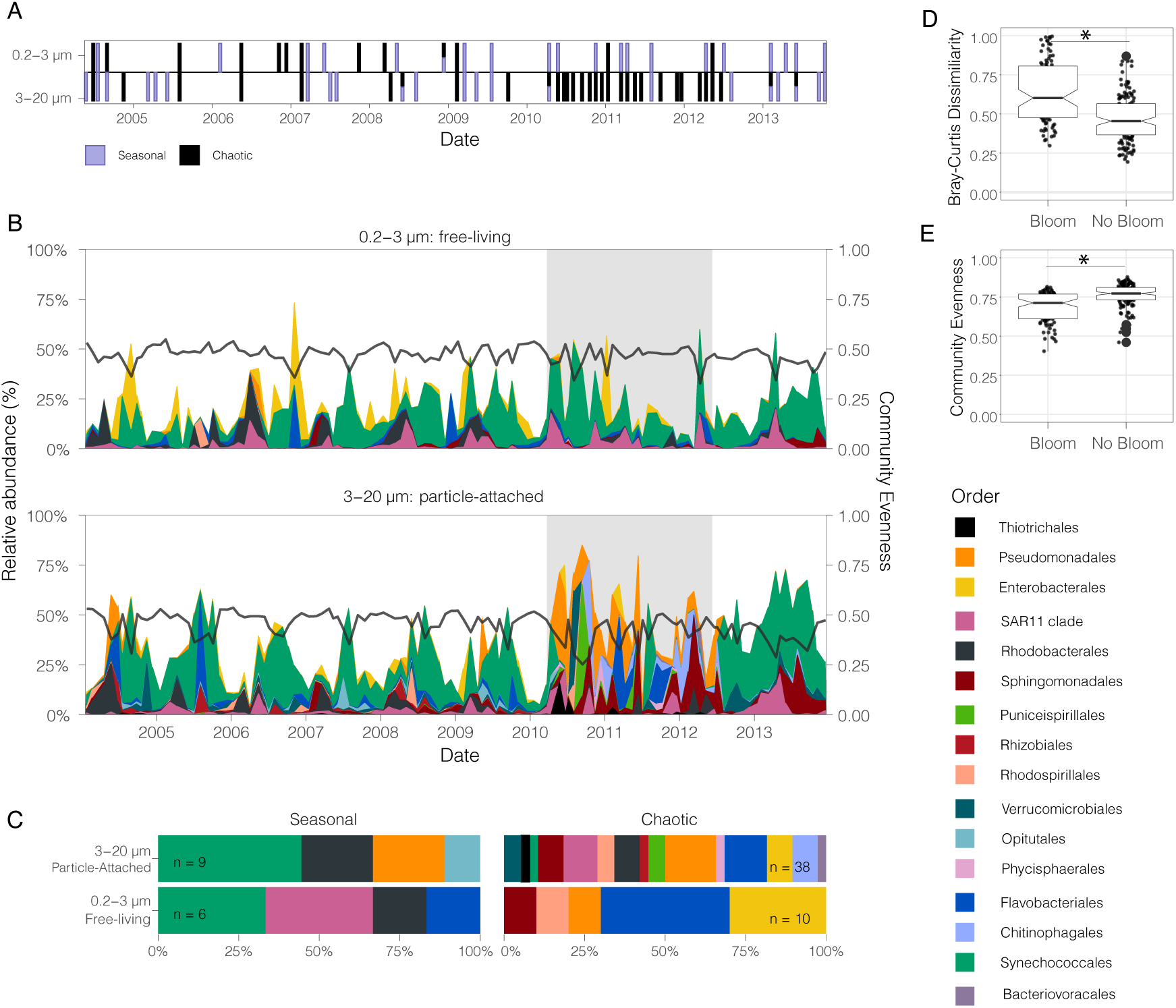
Abundance and taxonomy of BBMO bloomers over time. (A) Time series of bloom events, with bloomer type indicated by color as either seasonal or chaotic. The x-axis represents the temporal series, and the y-axis represents the size fraction. (B) Relative abundance (%) of bloomers across the 10-year time series at the Blanes Bay Microbial Observatory (BBMO), grouped by taxonomic order for the 0.2–3 µm and 3–20 µm size fractions. Colors represent different taxonomic orders, and the shaded area indicates the harbor restoration period. Solid lines indicate community evenness for each size fraction. (C) Taxonomic composition of the two main bloomer categories identified by wavelet analysis: seasonal (n ASVs exhibiting recurrence every 8–16 months) and chaotic (n ASVs showing no significant recurrence). (D) Boxplots of Bray–Curtis dissimilarity between consecutive samples during bloom and non-bloom events. (E) Boxplots of community evenness during bloom and non-bloom events. Asterisks indicate significant differences between groups.

Bloomers were categorized based on their recurrence (recurrent or chaotic) and their occurrence in the community (i.e. ‘broad’, ‘intermediate’, or ‘narrow’, see Methods). As the recurrence of the bloomers in the time series was mainly between 8 and 16 months (see Fig. S6B and Supplementary Information II for details), we refer to them as ‘seasonal’ (Fig. S6B, see Methods). Most of the blooming taxa belonged to the narrow chaotic category (N= 6 for the 0.2-3 μm fraction and N= 29 for the 3-20 μm size fraction), but 6 seasonal bloomers were detected in the 0.2-3 μm size fraction and 9 in the 3-20 μm size fraction (Fig. 2B, Table S3). The seasonal bloomers belonged mainly to the orders Synechococcales and Rhodobacterales in both size fractions (Fig 2B, Fig. S7). The chaotic bloomers belonged mainly to Enterobacterales and Flavobacteriales in the 0.2-3 μm fraction, and to Pseudomonadales in the 3-20 μm fraction. Notably, as we used a 10% threshold to define a bloom event, some of the seasonal bloomers did not bloom every year, despite their recurrent patterns (Fig. S8).

### Response of blooming taxa to environmental perturbations

The wavelet spectra of several blooming taxa experienced significant changes from April 2010 to June 2012 (Supplementary Information II). This period coincided with the restoration of the Blanes harbor located near the sampling site, with the most intensive work and consequent sediment resuspension taking place from April 2010 and June 2010 [45]. Accordingly, we observed notable alterations in the dynamics of the bloomers in the 3-20 μm fraction (Fig. 2A). Specifically, two-thirds of the blooming bacteria identified in this period (N= 32/47) showed significant shifts in abundance associated with the harbor restoration (Fig. S9). Overall, we were able to identify distinct responses among taxa i) seasonal taxa negatively affected by the perturbation, ii) chaotic taxa that disappeared during the restoration period, and iii) chaotic taxa that increased during that time (Fig. 3, Fig. S10). Moreover, the Shannon diversity index decreased significantly (p-value < 0.05) in the 3-20 μm size fraction during and after the restoration, while no significant change was observed in the 0.2-3 μm fraction (Fig. S11).

**Figure 3.**
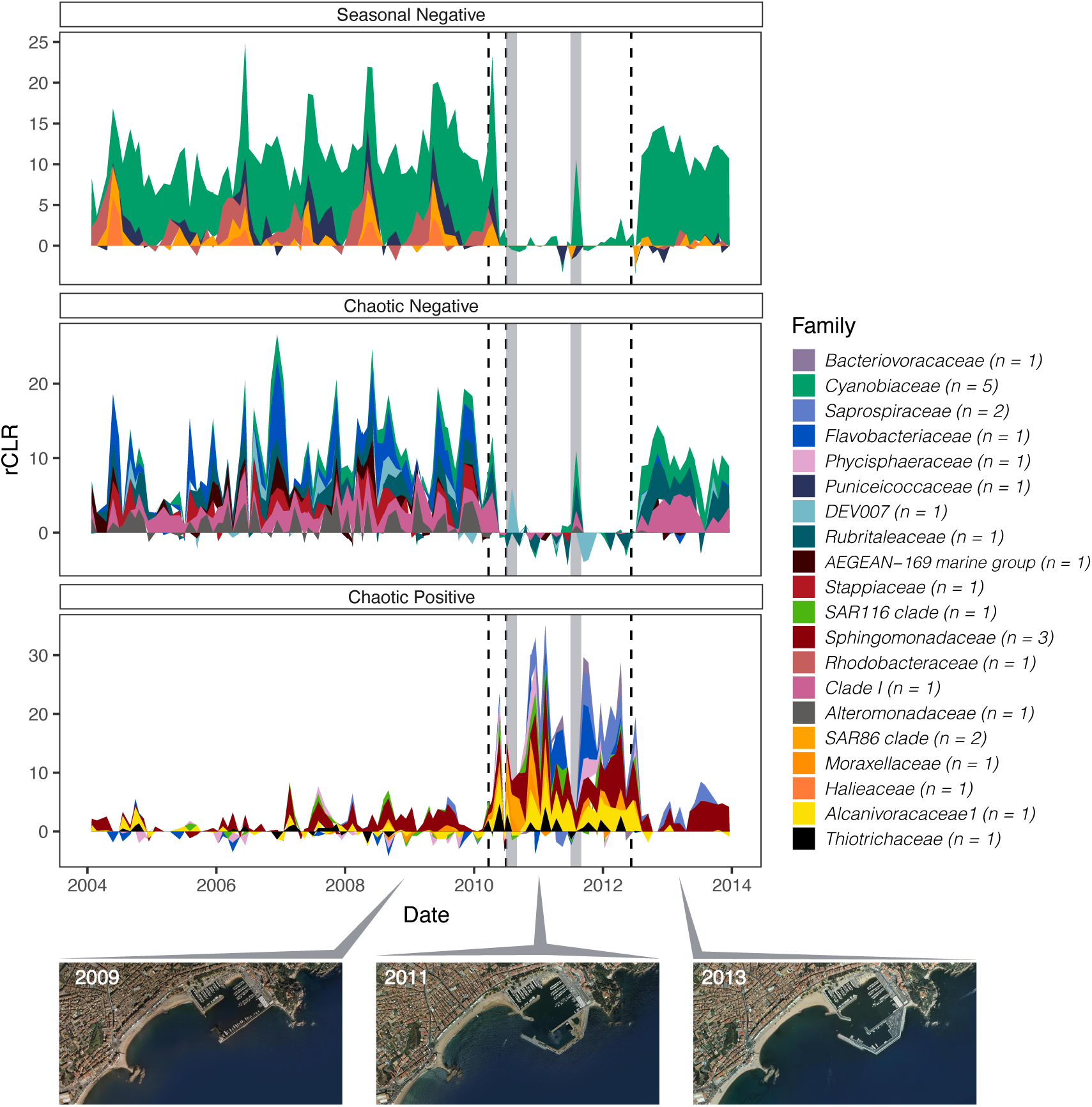
Response of the 3-20 µm size fraction blooming taxa to environmental perturbations. A) Robust centered log ratio (rCLR) abundances of those ASVs whose abundance values were significantly affected by the harbor restoration (according to one-way ANOVA or Kruskal-Wallis tests when normality was not met). Dashed lines represent the beginning of the harbor restoration (March, 2010), the end of the most important works (July, 2010), and the end of the harbor restoration (April, 2012). The grey shaded areas in the time series correspond to the summer period during harbor restoration, when the works stopped due to the high tourism season in the area. Upper panel: seasonal blooming ASVs negatively affected by the perturbation, central panel: chaotic blooming ASVs negatively affected by the perturbation, and lower panel: chaotic blooming ASVs positively affected by the perturbation. The n value indicates the number of ASVs included in each taxonomic family. At the bottom, orthoimages of the Blanes harbor before (2009), during (2011) and after (2013) the restoration, from Institut Cartogràfic i Geològic de Catalunya (ICGC), under the license CC BY 4.0.

### Low phylogenetic coherence in the blooming-trait

We explored whether blooming behavior is a taxonomically coherent trait, and whether bloomers were affected by the presence of closely related taxa. To do so, we analyzed the taxonomic diversity of the detected bloomers and examined their patterns of abundance in relation to their closest relatives. The bloomers detected in our dataset belonged to 6 different taxonomic phyla (Table S3). Around 20% of the bloomers (11 out of 57) had a phylogenetically closely related bloomer (i.e. defined as having less than 5 base pairs different in their ASVs sequences [14, 34]). In families such as SAR11 clade I and *Cyanobiaceae*, bloomers were generally more closely related to each other than to other ASVs classified as non-bloomers within the same family, whereas this pattern was not observed in the other families (Fig. S12). Seven of the bloomers with a close phylogenetic bloomer presented opposite patterns of abundance over the time series (Fig. 4A, S13A,C). In contrast, two closely related bloomers belonging to the genus *Synechococcus* exhibited synchronized dynamics over time until 2009, after which they began to decouple in the 0.2-3 µm size fraction (Fig 4B). In the family *Sphingomonadaceae,* the bloomer ASV17 occurred over the whole time series whereas ASV77 was only occasionally detected (Fig. 4C), yet they coexisted in two occasions. The rest of the bloomers had no closely related blooming ASVs and typically did not coexist with their closely related non-blooming ASVs (see Fig. S13E,F for an example).

**Figure 4.**
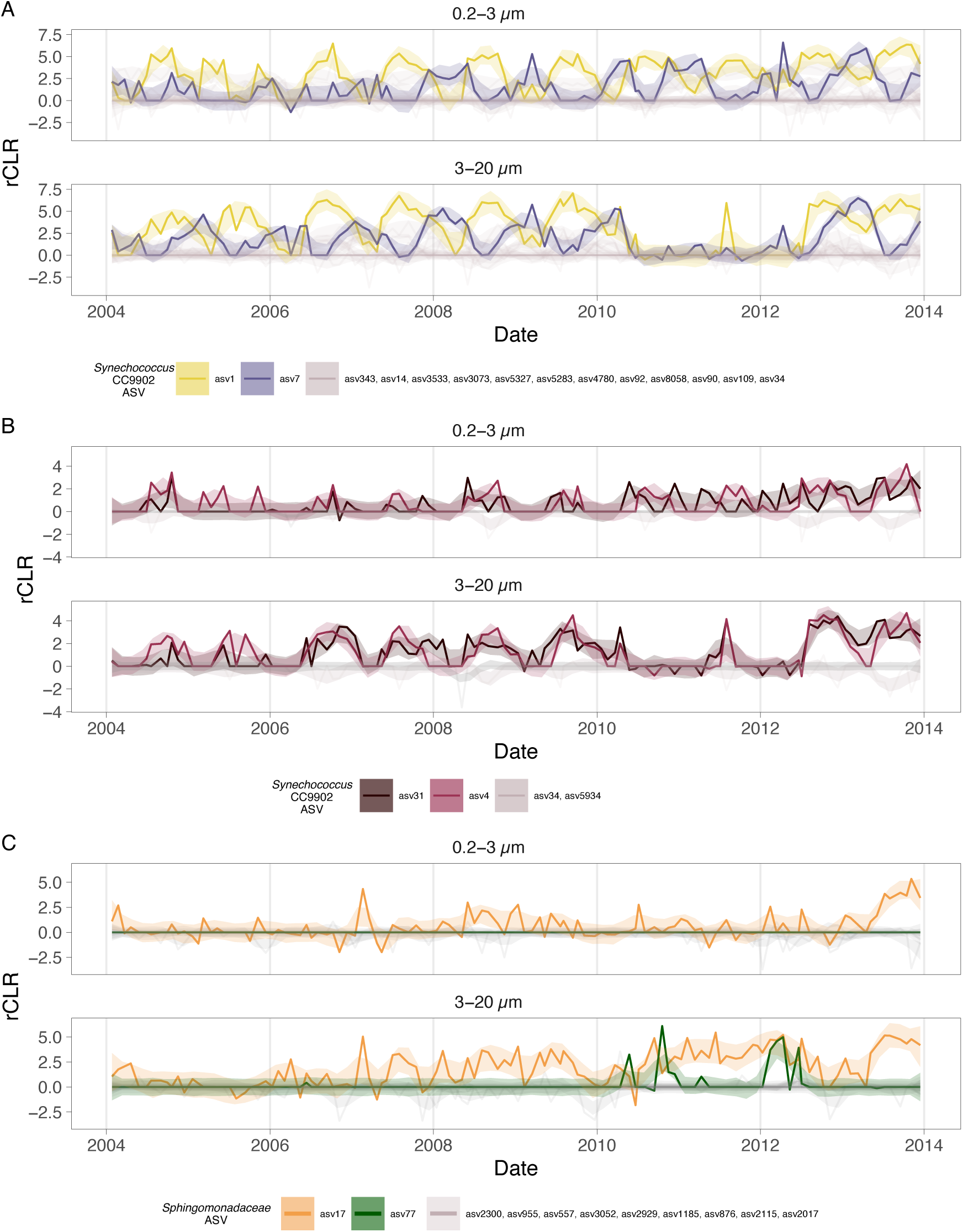
Time series of rCLR transformed abundances of representative bloomers (sharp colors) and their relationships with their closely related ASVs (p-distance < 0.012, which means less than 5 base pairs difference between ASVs, soft colors). Abundances are slightly smoothed for a better visualization of the results (LOESS span 0.086).

### Stronger causal relationships among pairs of ASVs in the 3-20 µm community

We further aimed to identify with CCM analyses causal relationships between broad-occurrence bloomers (5 ASVs in the 0.2-3 μm size fraction and 7 ASVs in the 3-20 μm size fraction) and the surrounding community, or the environmental variables. To summarize the results, we grouped the variables tested for causality into four categories: ASVs present in the 0.2-3 μm fraction, ASVs present in the 3-20 μm fraction, and biological or physicochemical environmental variables, and explored whether there were significant differences (for bloomers and non-bloomers) in the degree of causality (measured as the predictive skill, rho) by category (Fig. 5). In general, the patterns for bloomers and non-bloomers were similar, the physicochemical and biological variables had weaker causality than the community, and the 3-20 μm community had significantly stronger intra-causal relationships. This means that the 3-20 μm community was better predicted by its own composition (Fig. 5). From all the tested environmental variables, the physicochemical variables had higher causality than the biological ones, especially for the 3-20 μm community, in which phosphate stood out for having a high predictive power on the abundance of several bloomers (Fig. S14). According to the CCM analysis, in the 3-20 µm size fraction, some SAR11 ASVs (both bloomers and non-bloomers) were the ones with a stronger causal relationship with other ASVs (Fig. S14). Focusing on the blooming community, *Alcanivorax–*ASV22, with a chaotic behavior, was the bloomer with the highest average causal effect on other ASVs in the community (Fig. S15).

**Figure 5.**
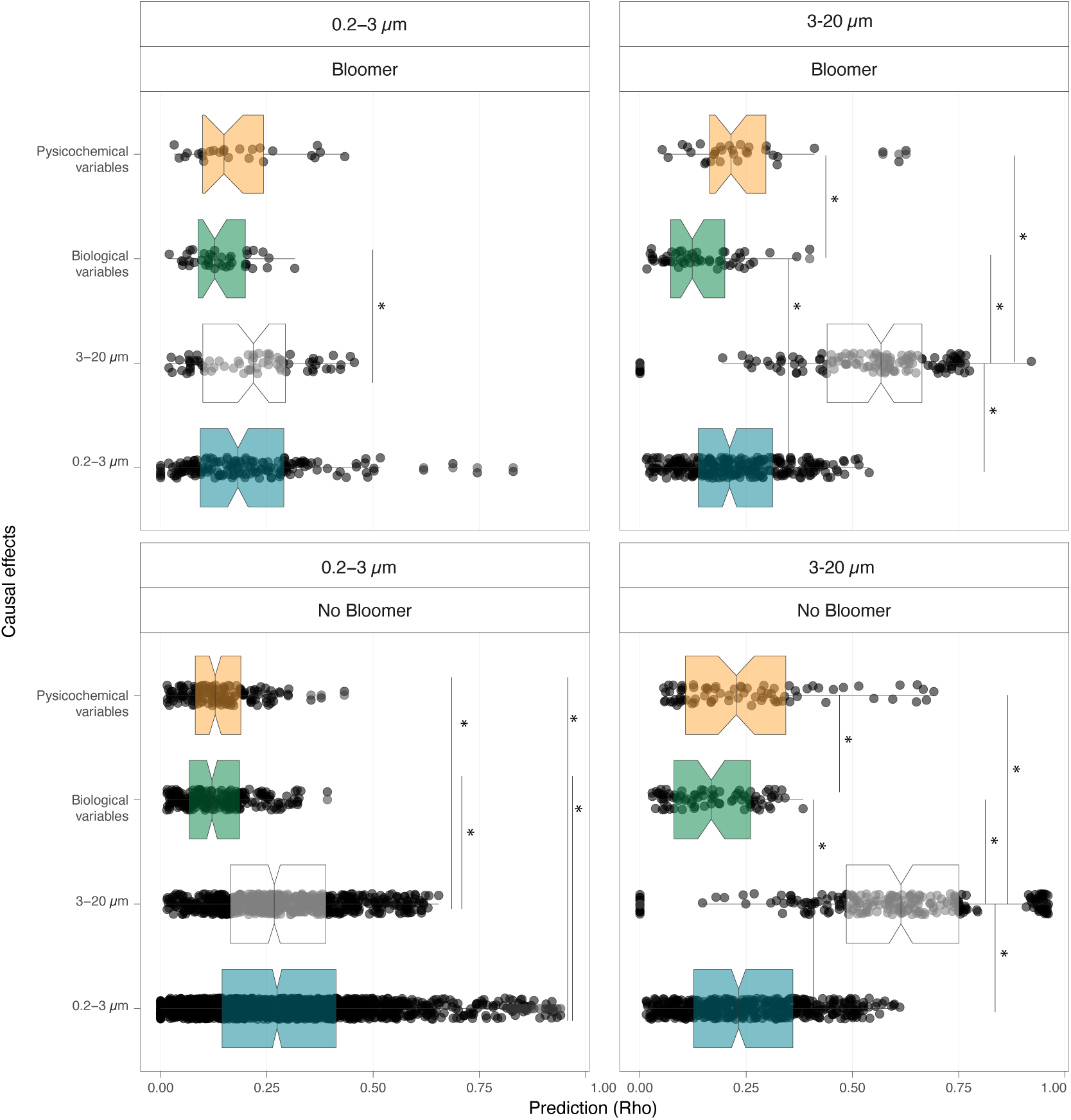
Comparison of the significant causal relationships of community composition (0.2-3 µm or 3-20 µm) and environmental variables (physicochemical and biological) on bloomer and non-bloomer ASVs based on Convergent Cross Mapping (CCM). The number of bloomers included in the analysis were 8, and non-bloomers 42. The Y-axis summarizes in boxplots the variables within each category that are significantly affecting the ASVs, categorized into two groups: bloomer (upper panel) and non-bloomer (second panel) in both size fractions, the 0.2-3 µm (first column) and 3-20 µm (second column). Statistically significant differences between groups are highlighted with an asterisk.

The CCM analysis also highlighted that *Synechococcus*–ASV7 and the SAR11 clade Ia–ASV15, exhibited a strong causal relationship over one another in both size fractions (Fig. S14). These ASVs were tightly coupled throughout the time series (Fig. 6A), co-occurring and displaying similar patterns across the entire observation period. Furthermore, the MDR S-map analysis, which allowed us to reconstruct the potential interaction networks over time, revealed that their interaction was stronger from *Synechococcus*–ASV7 towards SAR11 clade Ia–ASV15, particularly in the 3-20 µm size fraction. These interactions were mostly positive over time (Fig. 6B), whereas SAR11 clade Ia–ASV15 had lower interactions with weaker influence on *Synechococcus*–ASV7 (Fig. S16).

**Figure. 6.**
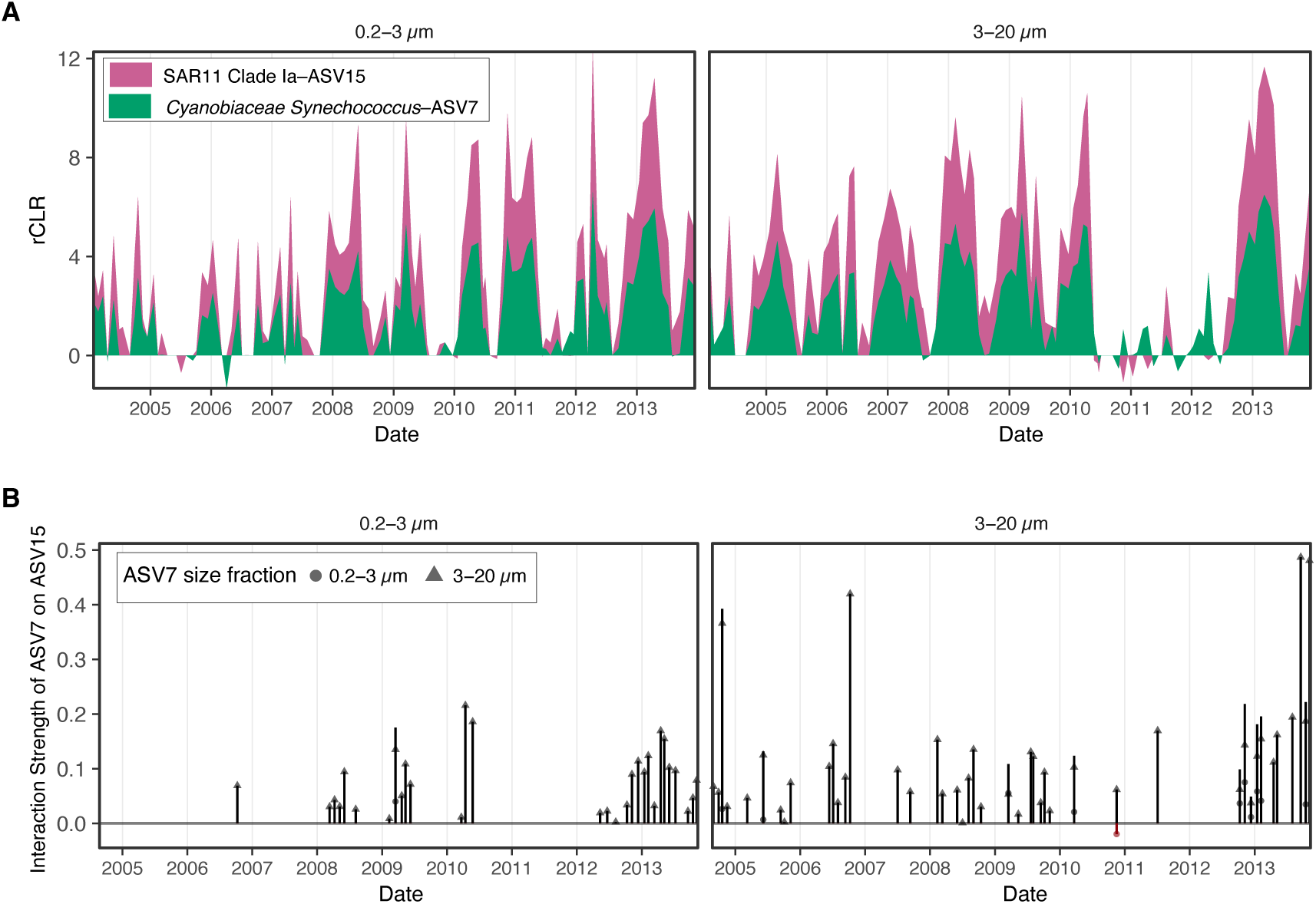
Example of co-existing taxa: A) Time series of the rCLR transformed abundances of ASV7 and ASV15, belonging to the *Synechococcus* and the SAR11 Clade Ia, respectively, that were co-occurring in the time series in both size fractions, and were strongly causally related. B) Interaction strength that ASV7 has on ASV15 during the time series according to Multiview distance regularized (MDR) S-map. Symbol shape indicates the size fraction from which ASV7 affects ASV15 in either the free-living size fraction (left panel) or the particle attached one (right panel).

### Blooming taxa altered the functional potential of the community

Finally, we explored whether the presence of a bloomer led to a change in community function using 5 years of metagenomic data from the 0.2-3 μm size fraction (years 2009-2013, see Methods). Beyond examining functional shifts, the comparison of metabarcoding and metagenomic datasets provides additional support that the observed blooming dynamics are not driven by PCR artifacts arising from the multiple rRNA operons typically present in bacterial copiotrophs. As mentioned above, blooming events were characterized by a significant shift in taxonomic composition (Fig. 2C). Consistently, the Bray-Curtis distance in gene composition between consecutive samplings presented significant correlations with taxonomic composition (N= 60, Spearman R= 0.73, p-value < 0.01) (Fig. 7A), confirming the blooming dynamics observed by metabarcoding. Then, we aimed to determine whether blooming events led to changes in community functionality, reflected in shifts in KOs composition. In this case, we also found a significant positive correlation between Bray-Curtis dissimilarity calculated on metabarcoding data and on KOs data (N= 60, Spearman, R: 0.64, p-value < 0.01). Nonetheless, the range of variation in the Bray-Curtis distance was much lower in the KO-based than in the gene-based composition. Focusing on the specific heterotrophic ASVs that bloomed during 2009-2014 in the 0.2-3 µm size fraction, we observed that the temporal patterns of gene abundances associated to the taxonomy of the bloomers exhibited consistent trends with the relative abundances of *Glaciecola*–ASV11 and *Amylibacter*–ASV27, whereas for the NS4 marine group–ASV58 the pattern was not as evident (Fig. 7C, D). Among the specific functions enriched during two particular bloom events, we thoroughly studied the top 25 KOs in which these bloomers were contributing the most (Table S4) and assessed their relative contribution to these functions within the context of the overall prokaryotic community. *Glaciecola*–*ASV11* (Fig. 7F) displayed a stronger contribution to several specific functions compared to *Amylibacter*–ASV27 (Fig. 7E). For instance, in January 2011, *Glaciecola*–*ASV11* sustained more than 75% of the functions associated to genes *smg* (K03747, conserved protein of unknown function, DUF494), *sapA* (K19226, an ABC transporter putatively involved in peptide transport), *aspC* (K00813 involved in amino acid transport and metabolism) and *pilE* (K05655, type IV pilus assembly protein). Overall, most of the functions associated to these bloomers were identified either as transporters, peptidases, or involved in carbohydrate and amino acid metabolism or DNA repair (Table S4).

**Figure 7.**
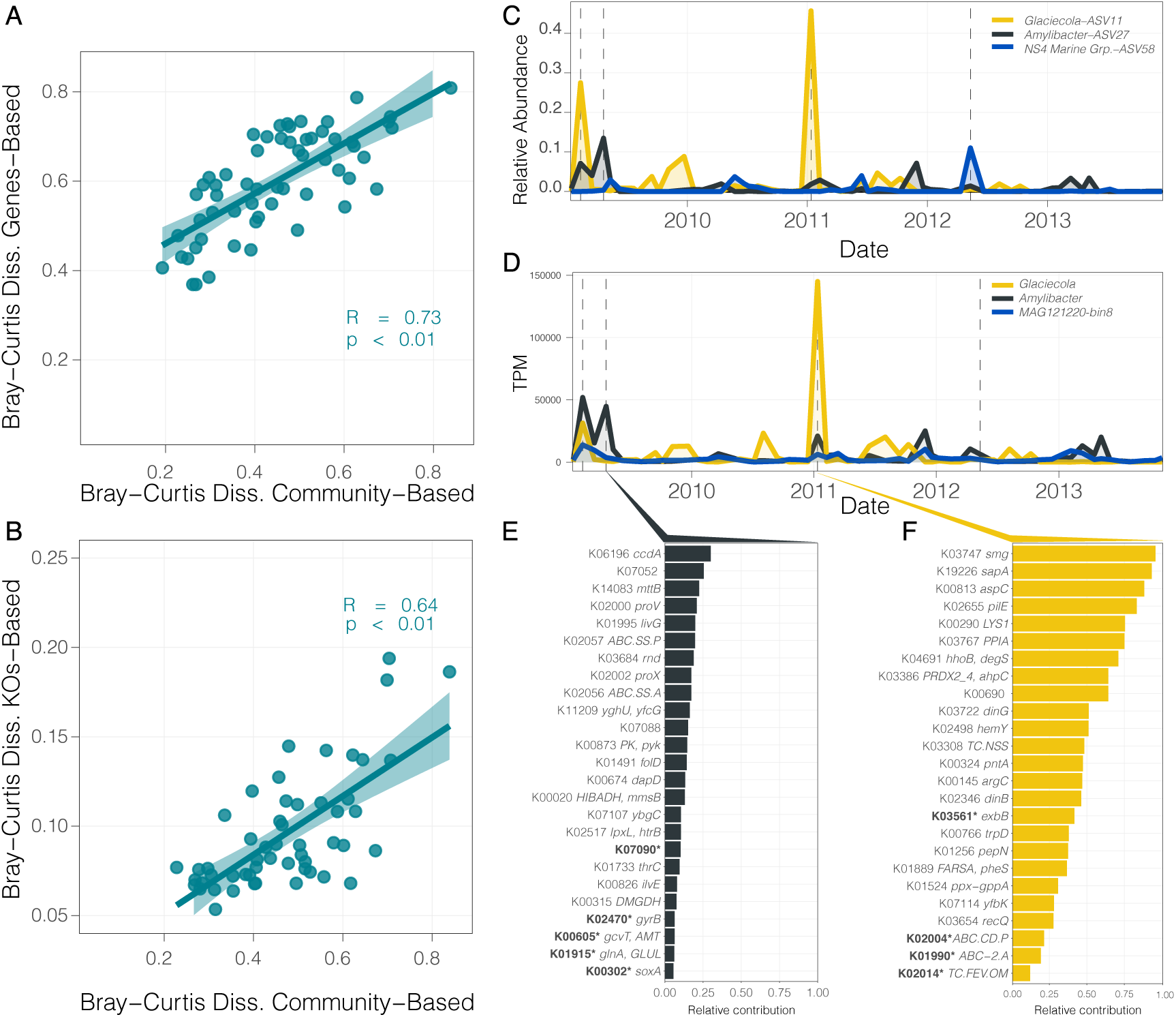
Potential functional implications of blooming events. Comparison of changes in ß-diversity metrics (i.e., Bray-Curtis dissimilarity based on community structure and genetic or functional composition) on consecutive samples of the 0.2-3 µm size fraction between 2009 and 2013 (A and B). Spearman correlation was used to assess correlations when normality was rejected (Shapiro test, p < 0.01). The correlations are as follows: A) Bray-Curtis dissimilarity of community composition versus Bray-Curtis based genetic content, B) Bray-Curtis community composition versus Bray-Curtis based on KEGG Orthologs (KOs) content. C) Relative abundance of heterotrophic bloomers between 2009 and 2014, where each line represents a specific ASV, denoted by a different color. D) TPM genes abundances transformed for all genes taxonomically identified within the bloomer taxa. E-F) Top 25 KOs sustained by specific bloomers during the bloom events of E) *Amylibacter* in April 2009, and F) *Glaciecola* in January 2011. The X-axis indicates the relative contribution of the bloomers to the total abundance of the KO. KOs highlighted with an asterisk and in bold indicate those that were also among the top 25 most abundant KOs in the whole community at the corresponding time point.

## Discussion

Previous studies have identified short-lived proliferations of specific prokaryotic taxa [23, 24], yet a detailed analysis of bacterial blooming behavior, the bloom frequency and their impact on community function have not been previously evaluated. Our work addresses this gap by integrating multi-year metabarcoding and metagenomic datasets, demonstrating that seasonal and chaotic bacterial blooms can be frequently detected in the marine environment even at a monthly sampling resolution. Among the seasonal and chaotic bloomers we found both phototrophic and heterotrophic ASVs. Seasonal bloomers were previously described seasonal taxa, such as members of Synechococcales [46], the SAR11 Clade [31, 47], or Rhodobacterales; the latter more linked to phytoplankton blooms [20, 48, 49]. Chaotic blooms were likewise frequent, but were driven by more opportunistic and copiotrophic groups, such as Flavobacteriales, Pseudomonadales or Enterobacterales [48, 50]. Their occurrence may be linked to pulses of organic matter [51], resistance to viral outbreaks [52] or other environmental perturbations, such as storms [53], or dust deposition events [54].

Despite the known strong seasonality and community succession at the BBMO [55, 56], we did not observe a significant correlation between consecutive changes in the ß-diversity and biological and physicochemical fluctuations (Fig. 1). This aligns with other studies that found that shifts in the environment were not consistently linearly linked to community changes [52, 57]. Instead, slight natural or human-induced changes have been proposed to trigger blooming events of specific taxa [7]. However, it is important to note that dynamic environmental fluctuations were likely not captured by our low sampling frequency, limiting the ability of the measured variables to explain bloom dynamics. Likewise, some key variables such as total organic carbon, or the quality of resources were not measured, limiting the identification of the triggers of bloom events. Nevertheless, our data show that biological interactions may also play a role in community dynamics (Fig. 5), in agreement with other studies [19, 58].

Knowledge on the dynamics of bloom-forming taxa in time series can help detect ecosystem changes and identify early indicators of environmental stress (e.g. meteorologic events) or anthropogenic perturbations (e.g. oil spills, wastewater discharges etc.). In strongly seasonal ecosystems such as the BBMO, community dynamics are largely driven by periodic environmental forcing [56]. Yet the remarkable shift in the blooming community from April 2010 to June 2012 likely reflects a disruption of the expected seasonal structure due to the Blanes harbor restoration works (Fig. 3). Interestingly, during the summer break of the restoration activities, we observed a reappearance of certain seasonal taxa that had been negatively affected by the restoration, indicating ecosystem resilience, only to disappear again once the works resumed (Fig. 3). The restoration activities were linked to increases in sediment resuspension, organic matter content in sediments, and deposition rates [45]. The bloomers detected in the 3–20 μm fraction could have been attached to these particles, but the 0.2-3 μm community was not altered by the sediment resuspension, perhaps because resuspended particles were large [45]. Still, the 0.2-3 μm community may have served as a reservoir of diversity or seed bank, facilitating recovery once prior conditions were restored [59]. An alternative explanation for the pronounced fluctuations in the 3-20 μm community during the restoration period is its stronger causal interconnectivity compared to the 0.2-3 μm community (Fig. 5, Fig S14), likely due to their coexistence on particles. This interdependence may increase susceptibility to cascading responses to external perturbations, making the 3–20 μm fraction a sensitive sentinel for ecosystem disturbances. Although the system recovered after harbor restoration, some bloomers in the 3-20 μm fraction that had dominated before the works did not regain their previous abundance levels even a year after the perturbation (Fig. S10). This suggests that some disturbances of the microbial community could have long-lasting effects. However, to fully assess ecosystem recovery and resilience, the analyses should be extended several years post-perturbation [60].

The ability to bloom may result from a combination of different traits, allowing taxa from diverse phyla to exhibit this capability. Most blooming ASVs lacked closely related taxa, consistent with the well-known competitive exclusion principle [61]. The ability to bloom may not be a taxonomically coherent trait, consistent with observations that closely related prokaryotes can differ in ecological behavior or in functional traits, such as lipid degradation capacities [62], growth rates [63] or environmental preferences [34, 64–66], leading to distinct patterns of abundance. However, our monthly sampling precludes firm conclusions about taxonomic coherence, as ASVs classified as non-bloomers may still bloom outside the temporal window captured by our study. Nevertheless, our results suggest that closely related blooming taxa can exhibit distinct ecological preferences, shaped by both environmental conditions and species interactions. An example of the complexity of these interactions can be observed in the blooming dynamics of SAR11 Clade Ia–ASV15 and *Synechococcus*–ASV7, which displayed tight co-occurrence patterns across both size fractions (Fig. 6). MDR analyses indicated a positive influence of ASV7 on ASV15, consistent with the previous observations that SAR11 at the BBMO is preferentially stimulated by *Synechococcus*-derived dissolved organic matter [67]. Similar metabolic dependencies have been reported between SAR11 and *Prochlorococcus* [68, 69]. Among the CCM-inferred causal relationships, within the broad bloomers, the chaotic *Alcanivora*x–ASV22, exerted the strongest influence on other ASVs (Fig. S15). Given the known oil-degrading ability of this genus [70], and its proliferation during the harbor restoration, this taxon may have played a key role in structuring community interactions under disturbed conditions.

The large changes in taxonomic composition typically associated to bloom events, led to functional shifts in the community (Fig. 7B), indicating low functional redundancy with the preceding community [71]. Despite the significant positive relationship, changes in the functional composition were smaller than those observed in community composition, particularly for KOs, likely due to shared functions across all living cells [72]. However, it is important to note that the lower changes in KO-based community composition may partly reflect annotation biases, as current databases capture only a subset of functional diversity with many genes not assigned to orthologous groups. Likewise, regarding the three heterotrophic bloomers detected, the accurate assignment of genes to NS4 Marine Group-ASV58 may have been hindered by low representation of this taxon in reference databases, whereas the trends for ASV11–*Glaciecola* and ASV27–*Amylibacter* could be clearly identified, mirroring the patterns observed through metabarcoding. Our results suggest that during bloom events these taxa could be disproportionally contributing to specific functions in the ecosystem. This was particularly evident for *ASV11– Glaciecola* (Fig. 7F), which alone sustained functions linked to genes such as *smg*, *sap*A*, pil*E and *aspC*. The enrichment in peptidases, transporters and genes involved in amino acid and carbohydrate metabolism (Table S4) further supports a strong role of blooming taxa in driving key microbial processes. Interestingly, similar functional enrichments have been reported in genomes of blooming taxa from mesocosm experiments [73]. The changes in abundance of bacterial bloomers could also be related to eukaryotic plankton abundances, for example, we found that the ASV27–*Amylibacter* showed high prevalence in interaction networks with several protists[74], potentially reflecting the utilization of phytoplankton exudates [21].

Altogether, this study provides the first long-term, detailed characterization of bacterial blooms in marine prokaryotic communities, showing that they are more frequent than hitherto thought. We show that bloomers strongly influence community composition, respond to anthropogenic disturbances and have an impact in the functional potential of the community. Our monthly time series sheds light on the frequency and recurrence of these events, but key aspects of bloom dynamics, including onset, duration and magnitude of the event, and the factors triggering the blooms, remain unresolved due to the limited temporal resolution. The pronounced chaotic behavior observed suggests that higher-frequency sampling will be necessary to unveil underlying mechanisms for bloom formation and the broader impact of these events in the marine ecosystem.

## Acknowledgements

We thank all those involved in maintaining the BBMO time series, particularly Clara Cardelús and Vanessa Balagué; Ramon Massana and Irene Forn for the flagellate data; and Lídia Montiel for metagenomic processing. A particular thought for our captain Anselm, who left us recently. We thank the Plankton Ecology Group at EAWAG for constructive feedback during the analyses, Guillem Roca, Jordi Pagès, and Teresa Alcoverro for insights on the harbor restoration period, and Ignasi Bartomeus for developing the bloomers’ function. This research was supported by grants RTI2018-101025-B-100, PID2021-125469NB-C31, and PID2022-136281NB-I00, PID2022-143213NB-100 from the Spanish Ministry of Science and Innovation. Ona Deulofeu-Capo was supported by a Spanish FPI grant and contract (MIAU: RTI2018-101025-B-100; PID2021-125469NB-C32). This work also acknowledges the Severo Ochoa Centre of Excellence accreditation (CEX2019-000928-S).

## Authors contributions

Data acquisition and preprocessing: IF, AA, XRV, RL. Conceptualization: ODC, MS, JMG. Analysis: CGC, ODC. Visualization: ODC. Supervision: MS, JMG, OS. Writing: ODC, CGC, MS, JMG. Funding acquisition: JMG, IF, EG, RL, OS. All authors contributed to the manuscript.

## Data Availability Statement

The raw metabarcoding data (PRJEB38773) and raw metagenomic data (PRJEB48035) are available in the European Nucleotide Archive. Genes tables are available in: https://zenodo.org/records/17183573, while the R-scripts used are available at: https://github.com/onadeulofeu/Bloomers_10y_BBMO and in https://github.com/EcologyR/Bloomers

