## Supplementary Information I for "The natural history of bacterial bloomers in a decade-long time series"

---

Index

**Table S1** (as a separate Excel file): Summary of physicochemical and biological variables measured during monthly sampling at the Blanes Bay Microbial Observatory. ______ 5

**Table S2** (as a separate Excel file): Summary of the identified blooming events across the BBMO-10Y timeseries and the ASVs that determined those events.______ 5

**Table S3** (as a separate Excel file): Summary of the potential bloomers identified in our time series, their occurrence across the time series, their recurrence and their taxonomy.______ 5

**Table S4** (as a separate Excel file): Details on the top 25 KEGG ortholog (KO) annotations that were most contributed by specific blooming taxa during two bloom events ______ 5

*Supplementary Figures:* ............................................................. *6*

**Figure S1:** Sensitivity analysis: exploration of different threshold values used to determine a blooming event in our dataset. ______ 6

**Figure S2:** Continuous wavelet analysis of four different ASVs in the 3-20 µm fraction, ASV84 (A), ASV311 (B), ASV85 (C) and ASV25 (D). ______ 7

**Figure S3:** Time series of smoothed total heterotrophic bacteria abundances and chlorophyll *a* concentration over the years.______ 8

**Figure S4:** Differences between the relative abundances of bloomers at each timepoint and the geometric mean of that ASV in the 0.2-3 µm size fraction. ______ 9

**Figure S5:** Differences between the relative abundances of bloomers at each timepoint and the geometric mean of that ASV in the 3-20 µm size fraction.______ 10

**Figure S6:** Categorization of ASVs based on their recurrence and their occurrence throughout the time series as determined using wavelets analyses.______ 12

**Figure S7**: Blooming community in our time series in each size fraction (0.2-3 µm or 3-20 µm) and whether they presented seasonal or chaotic fluctuations over the years.______ 13

**Figure S8:** Seasonal bloomers and their interannual variability. ______ 14

**Figure S9:** Statistical differences before, during and after the harbor restoration of bloomer ASVs abundances in the 3-20 µm size fraction (n = 47).______ 15

**Figure S10:** Relative abundance (%) of the different groups of ASVs affected by the harbor restoration in the 3-20 µm size fraction. ______ 16

**Figure S11:** Bacterial diversity changes during harbor restoration. ______ 17

**Figure S12:** Boxplots comparing pairwise phylogenetic distances between blooming and non-blooming ASVs, and among blooming ASVs, across taxonomic families containing more than one blooming taxon. Groups without boxplots reflect insufficient numbers of pairwise comparisons. Significant differences were assessed using a non-parametric Wilcoxon test and are highlighted with an asterisk. The color of the boxplots indicates the phylogenetic Order. ____ 18

**Figure S13:** Examples of the time-series of closely related bloomers (in bright colors) and their closely related ASVs (p-distance < 0.012, in dimmed colors) in the SAR11 clade (A), SAR86 Clade (C) and genus *Glaciecola* (E). ____ 19

**Figure S14:** Convergent Cross Mapping (CCM) results showing the causality of different ASVs and environmental variables on the bloomers. ____ 20

**Figure S15:** Boxplot of occurrent ASVs (> 1/3 occurrence) ordered by their causal effect on other ASVs in the community, identified using Convergent Cross Mapping (CCM). ____ 21

**Figure S16:** Multiview distance regularized (MDR) S-map presenting the interaction strength that SAR 11 Clade Ia–ASV15 has on *Synechococcus*–ASV7 during the time series. ____ 22

### Supplementary Methods

*Physicochemical and biological variables collected*

The following physicochemical parameters were measured: water temperature (ºC) and salinity (obtained in situ with a SAIV-AS-SD204 Conductivity-Temperature-Depth probe), day-length (hours of light), transparency (Secchi depth, in m), and the concentration of five inorganic nutrients ($PO_4^{3-}$, $NH_4^+$, $NO_2^-$, $NO_3^-$ and $SiO_4^{4-}$ determined with an Alliance Evolution II autoanalyzer). Abundances of heterotrophic bacteria, photosynthetic picophytoplankton and viruses were determined by flow cytometry using a FACSCalibur (BectonDickinson) flow cytometer [1]. To enumerate heterotrophic nanoflagellates, Cryptomonads, and Micromonas-like flagellates, seawater was filtered onto polycarbonate 0.6-µm filters, stained with 4',6'-diaminidino-2-phenylin-dole (DAPI, final concentration 1 µg·mL-1 [2], and counted in an Olympus BX61epifluorescence microscope. Total and <3 µm chlorophyll-a concentration was estimated through fluorometry of acetone extracts after filtration of 150 mL of seawater on GF/F filters. Bacterial production was estimated from the rates of protein synthesis using tritiated leucine [3].

*DNA extraction and sequencing*

DNA was extracted as described in [4] and stored at -80ºC. The metabarcoding sequencing was performed using a MiSeq sequencer (2 × 250 bp, Illumina) at the Research and Testing Laboratory (Lubbock, TX, USA; using the bacterial primers 341F (5′-CCTACGGGNGGCWGCA G-3′) [5] and 806RB (5′-GGACTACNVGGGTWTCTAAT-3′) [6] which amplify the V3–V4 regions of the 16S rRNA gene. There were three missing samples from the 3-20 µm size fraction for February and May 2004, and March 2005. Bioinformatic processing was performed at the Marine Bioinformatics platform (MARBITS: https://marbits.icm.csic.es/). Primers were trimmed using cutadapt v.1.16 [7]. DADA2 [8] was used to generate V3-V4 16S rRNA gene amplicon sequence variants (ASVs), and the taxonomic assignment was performed using the SILVA database v.138.1 [9] as reference.

*Metagenome processing*

Metagenomes were assembled individually into contigs with Megahit [10] and protein coding regions were predicted within the contigs using Prodigal v2.6.3 [11]. Genes were clustered at 95% identity and 80% coverage using Linclust v10 [12] to create a catalogue with non-redundant genes. Genes were taxonomically annotated using MMSEQ2 v11 -e1a1c [13] against Genome Taxonomy Database (GTBD:r89) [14]. To obtain gene abundances, reads were mapped to the gene catalog and normalized by gene length. We then applied two additional layers of normalization. First, to estimate the relative contribution of different bloomers to the community metagenome —comparable to relative abundance from 16S metabarcoding data—, gene counts were normalized to total per million (TPM) by sequencing depth. Second, to assess changes in genetic or Kyoto Encyclopedia of Genes and Genomes (KEGG) composition and quantify bloomers' contributions to specific KEGG Orthologs (KOs), counts were normalized using the geometric mean of the log-transformed counts of 10 universal single-copy genes (SCGs) using the COG functional annotation via rpsblast v2.7.1 that can be extrapolated to the respective KOs numbers in brakets: COG0012 *YchF*, GTP1 Ribosome-binding ATPase, GTP1/OBG family (K06942), COG0016 *PheS*; Phenylalanyl-tRNA synthetase alpha subunit (K01889), COG0018 *ArgS*; Arginyl-tRNA synthetase (K01887), COG0172 *SerS*; Seryl-tRNA synthetase (K01875), COG0215 CysS; Cysteinyl-tRNA synthetase (K01883), COG0495 *LeuS*; Leucyl-tRNA synthetase (K01869), COG0525 *ValS*; Valyl-tRNA synthetase (K01873), COG0533 TsaD; tRNA A37 threonylcarbamoyltransferase TsaD (K01409), COG0541 *Ffh*; Signal recognition particle GTPase (K03106) and COG0552 *FtsY*; Signal recognition particle GTPase (K03110) [15–17]. Genes were functionally annotated using the KEGG database (r20170220) via DIAMOND v0.9.22.

#### Supplementary Tables

**Table S1** (as a separate Excel file): Summary of physicochemical and biological variables measured during monthly sampling at the Blanes Bay Microbial Observatory.

**Table S2** (as a separate Excel file): Summary of the identified blooming events across the BBMO-10Y timeseries and the ASVs that determined those events.

**Table S3** (as a separate Excel file): Summary of the potential bloomers identified in our time series, their occurrence across the time series, their recurrence and their taxonomy.

**Table S4** (as a separate Excel file): Details on the top 25 KEGG ortholog (KO) annotations that were most contributed by specific blooming taxa during two bloom events (Figure 7), based on the Kyoto Encyclopedia of Genes and Genomes (KEGG) and its higher-order classifications (BRITE).

#### Supplementary Figures:

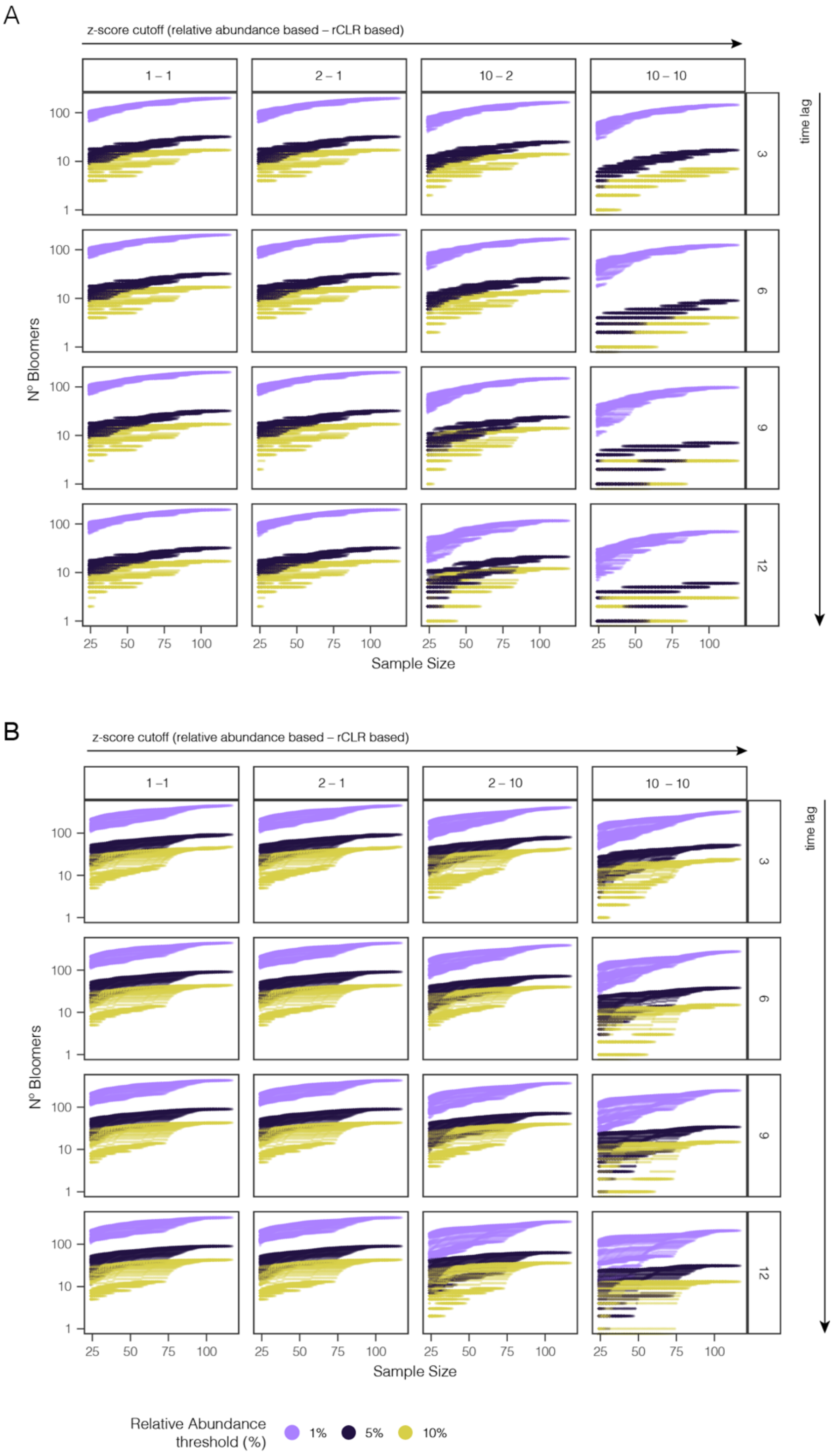

**Figure S1:** Sensitivity analysis of bloomers thresholds. Number of ASVs classified as bloomers under different threshold combinations: (i) minimum number of samples considered (x-axis), (ii) z-score thresholds calculated from relative abundance and rCLR values (columns), where the value before the hyphen corresponds to the relative abundance threshold and the value after the hyphen corresponds to the rCLR threshold (e.g., 2–2), and (iii) minimum number of samples required to compute z-scores (rows). Panel A corresponds to the 0.2–3 µm size fraction, whereas Panel B corresponds to the 3–20 µm size fraction. Note that axis Y is in log scale.

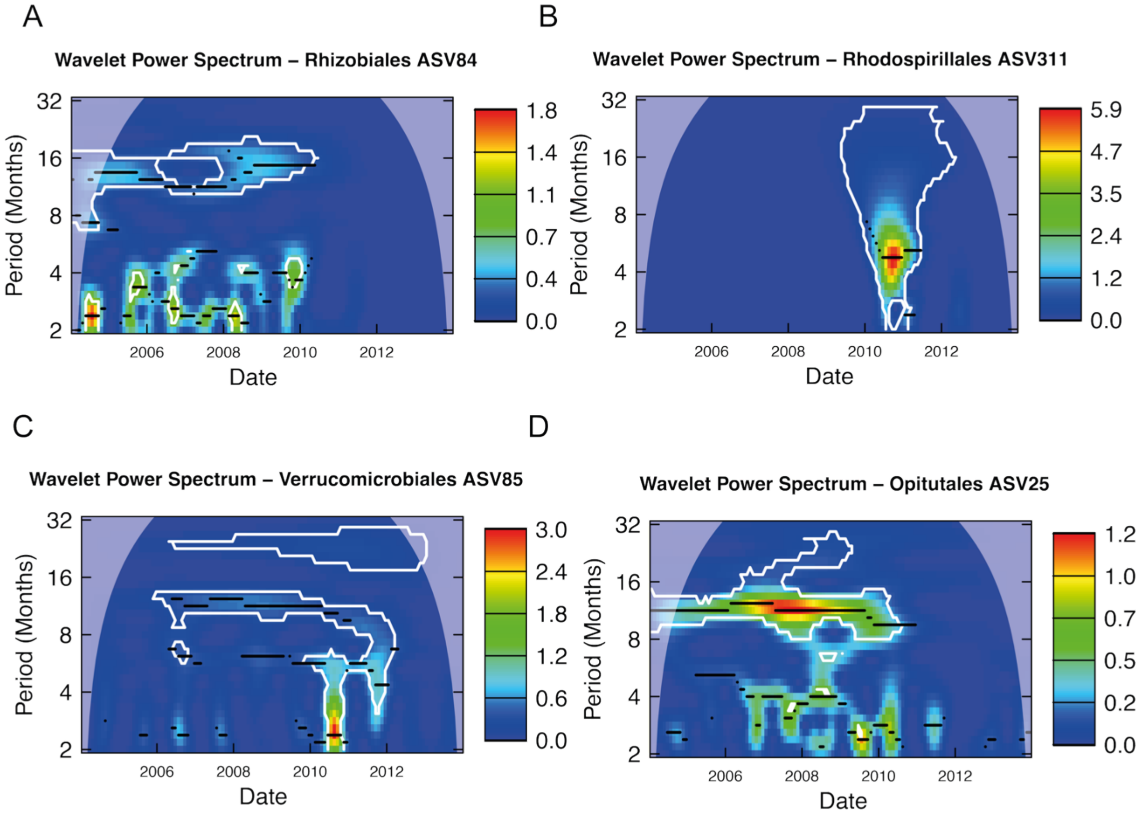

**Figure S2:** Continuous wavelet analysis of four different ASVs in the 3-20 μm fraction, ASV84 (A), ASV311 (B), ASV85 (C) and ASV25 (D). The X-axis represents the time series (2004-2014), and the Y-axis shows the period (in months). Color intensity indicates wavelet power, reflecting the strength of periodic signals. Significant (p-value after consistent results with 100 repetitions) signals within a period are highlighted by contour lines, while the color scale suggests that a periodic phenomenon is detected in the pattern of abundance of each ASV in that period. Stronger signal intensities at specific periods suggest recurrent patterns in ASV abundance. Temporal scales are interpreted as: short-term (2–4 months), semiannual (4–8 months), seasonal (8–16 months), annual (16–32 months), and interannual (~32 months). In case that an ASV has a recurrent pattern in one period we would observe a higher signal intensity at that period. In these examples, ASV84 (A) uncovers a significant annual periodicity (that oscillates between 8-16 months) and a stronger high frequency (2-4 months) periodicity that varies from year to year. Both disappearing in the last part of the time series. ASV311 (D), shows the highest power spectrum of these four examples, indicating strong periodic signals (~ 6 months) during the years 2010-2011.

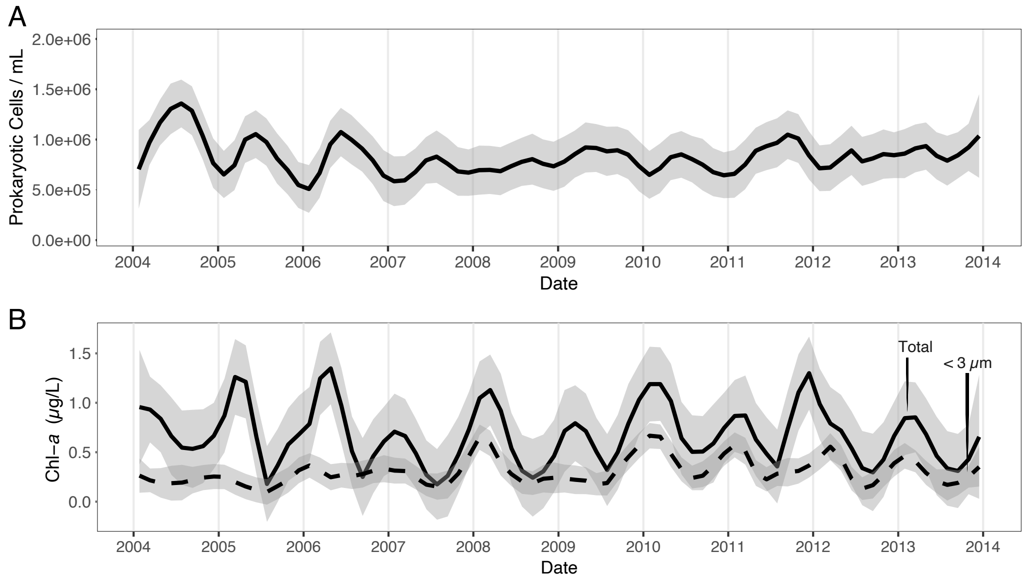

**Figure S3:** Time series of smoothed total heterotrophic bacteria abundances and chlorophyll *a* concentration over the years. A) Prokaryotic abundance values (cell/mL). B. Chlorophyll *a* (Chl-*a*) in µg/L. Dashed line is for the < 3 µm fraction, and the solid one is for the total chlorophyll *a* concentration. The shaded area represents the confidence interval around the smooth.

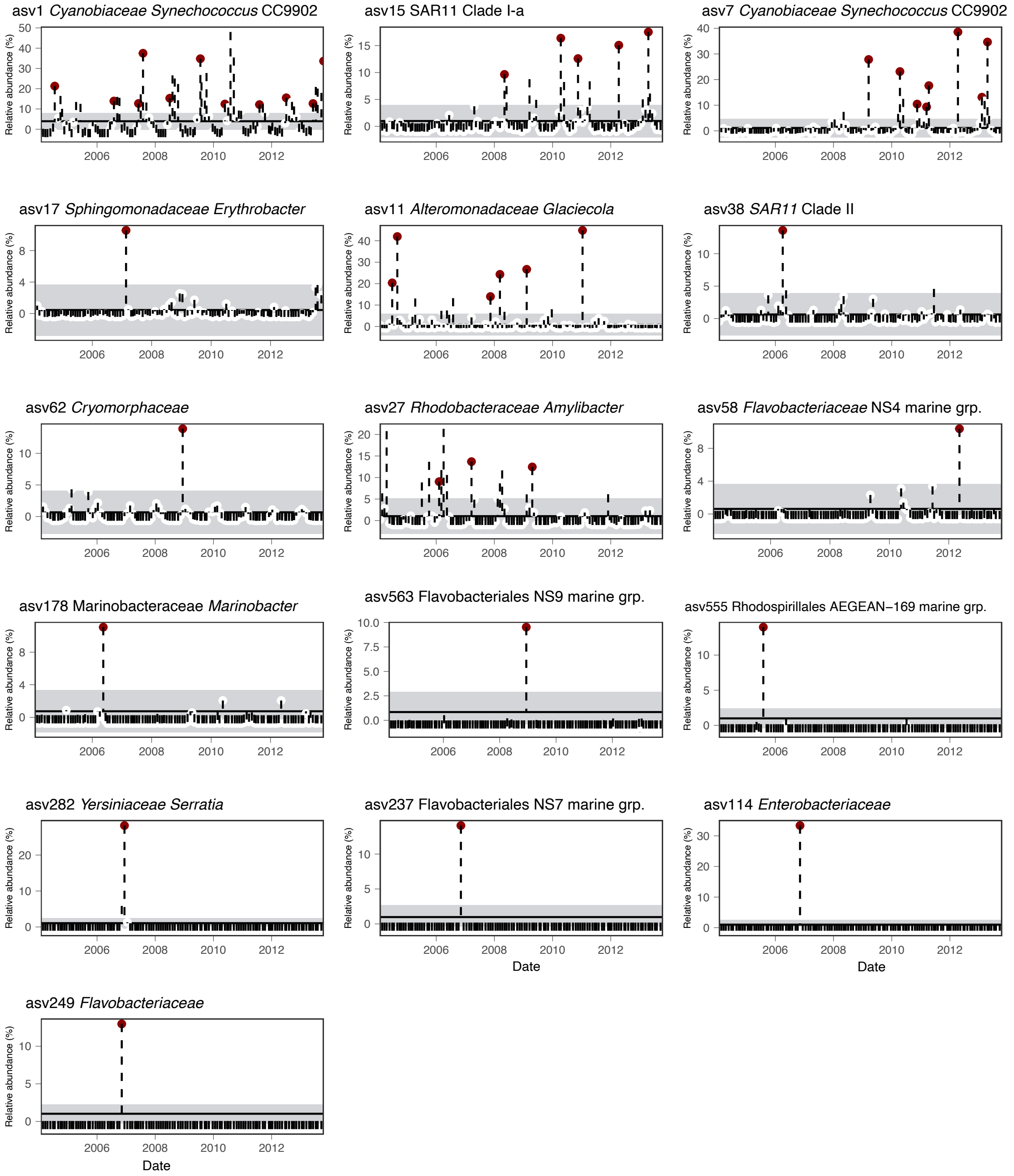

**Figure S4:** Differences between the relative abundances of bloomers at each timepoint and the geometric mean of that ASV in the 0.2-3 µm size fraction. Points are datapoints classified as blooming events (relative abundance > 10% and z-score > 2 based on relative abundance and > 1 based on transformed rCLR abundances) in our time series. The shaded area is the geometric standard deviation of the geometric mean for each ASV throughout the whole time series. The X-axis represents the different timepoints and the Y-axis the relative abundance of the sequence/ASV/organism in the community (%).

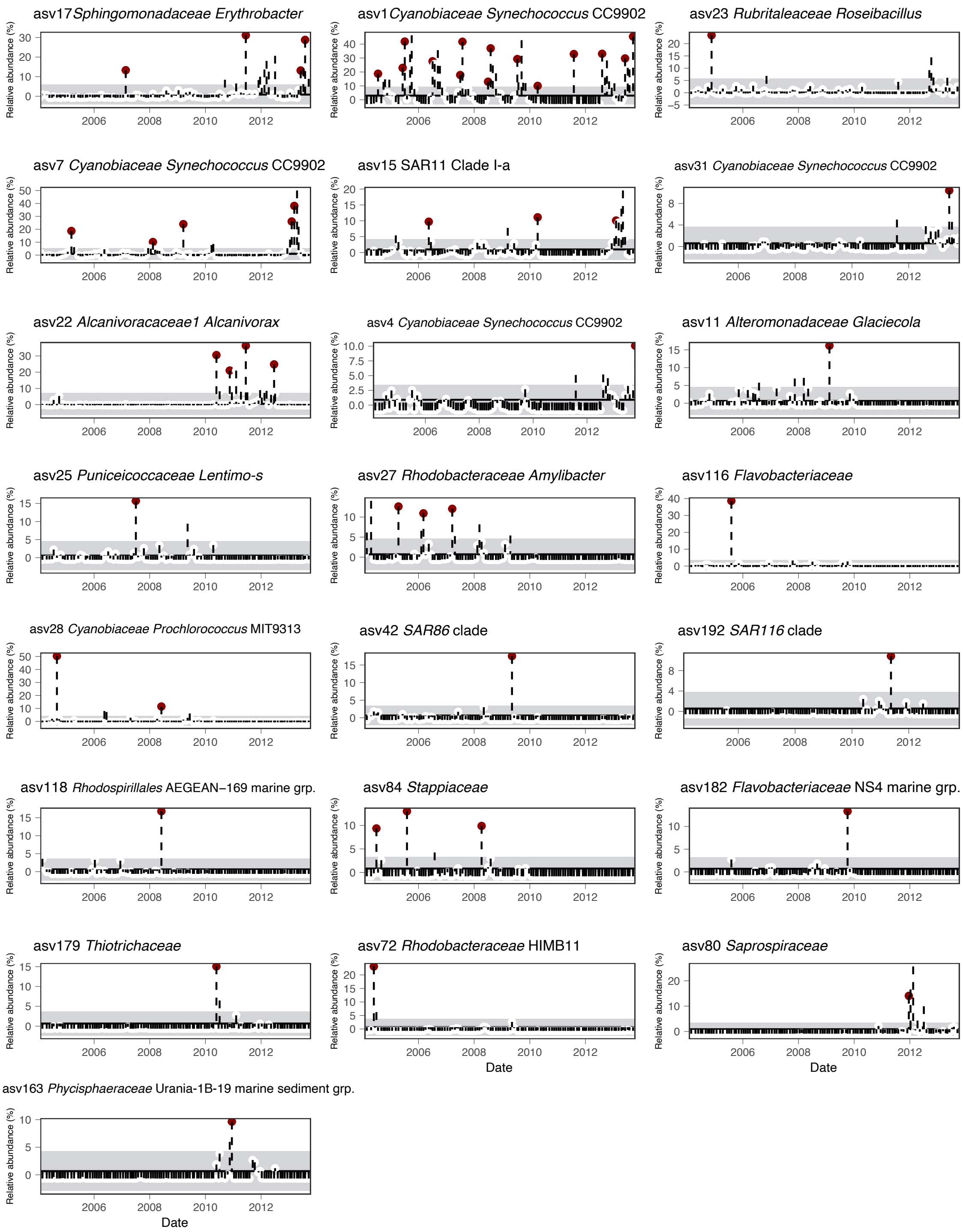

**Figure S5:** Differences between the relative abundances of bloomers at each timepoint and the geometric mean of that ASV in the 3-20 µm size fraction. Points are datapoints classified as blooming events (relative abundance > 10% and z-score > 2 based on relative abundance and > 1 based on transformed rCLR abundances) in our time series. The shaded area is the geometric standard deviation of the geometric mean for each ASV throughout the whole time series. The X-axis represents the different timepoints and the Y-axis the relative abundance in the community (%).

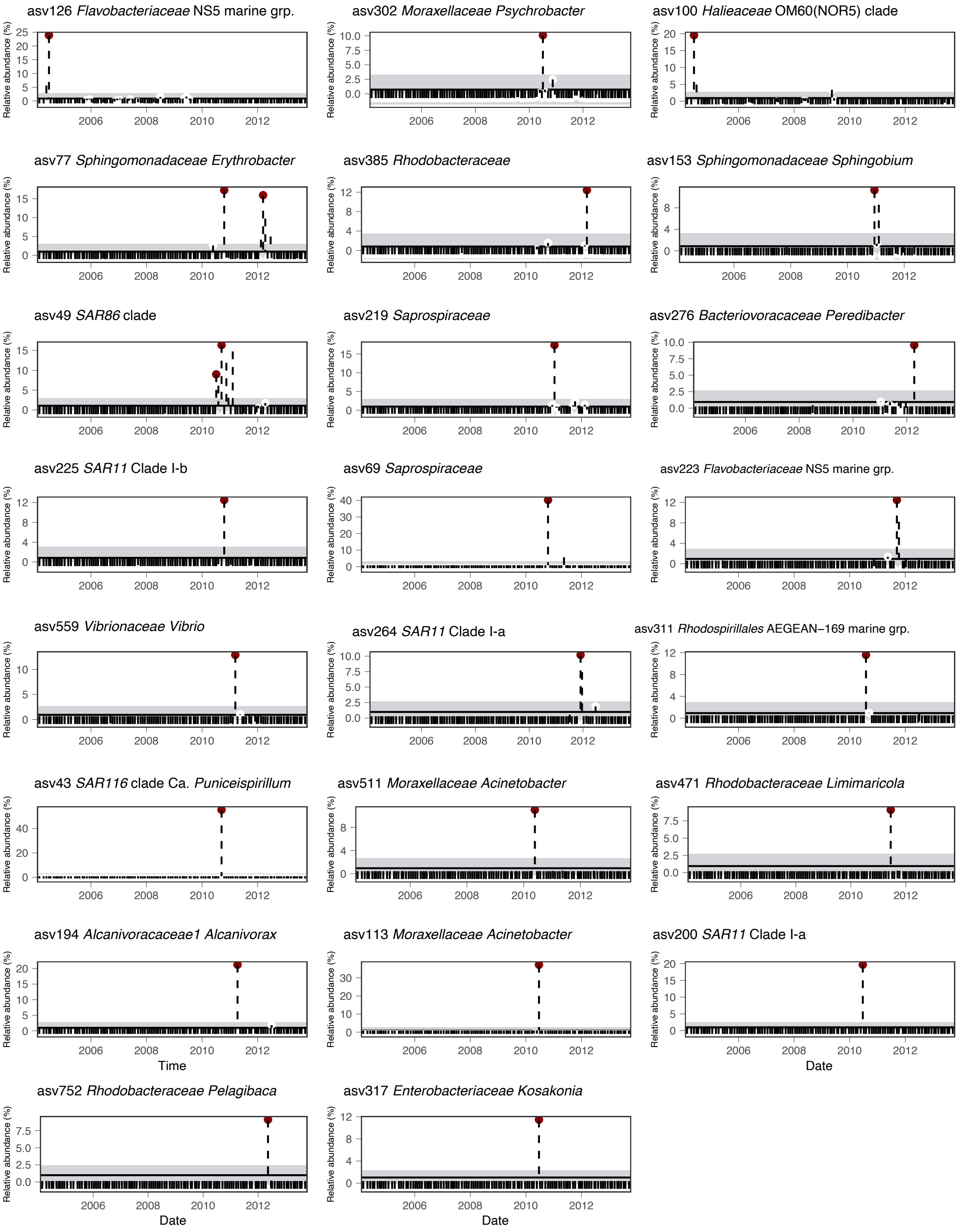

**Figure S5** (continuation): Differences between the relative abundances of apparent bloomers at each timepoint and the geometric mean of that ASV in the 3-20 μm size fraction. Points are datapoints classified as blooming events (relative abundance > 10% and z-score > 2 based on relative abundance and > 1 based on transformed rCLR abundances) in our time series. The shaded area is the geometric standard deviation of the geometric mean for each ASV throughout the whole time series. The X-axis represents the different timepoints and the Y-axis the relative abundance in the community (%).

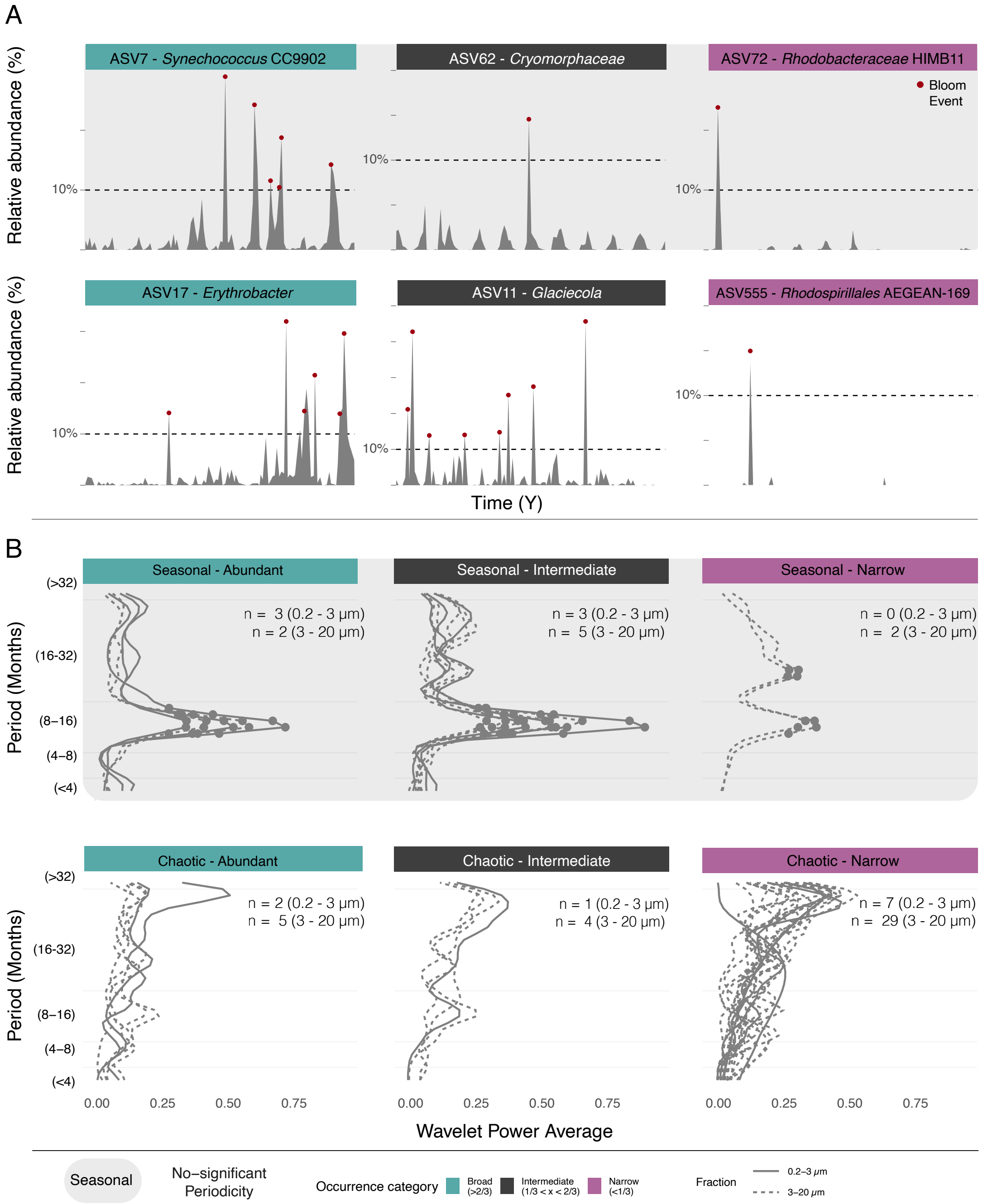

**Figure S6:** Categorization of ASVs based on their recurrence and their occurrence throughout the time series as determined using wavelets analyses. A) Temporal dynamics of 6 different blooming ASVs that exemplify the different types of bloomers detected at the BBMO. The dashed line indicates the 10% of the community as threshold for a blooming event. The dots indicate blooming events based on the z-score and the relative abundance threshold. In the first row, three examples of recurrent bloomers (ASV7 (0.2-3 μm), ASV62 (0.2-3 μm), ASV72 (3-20 μm)) based on the wavelets analysis. In the second row there are those bloomers which were not recurrent (i.e. chaotic) (ASV17 (3-20 μm), ASV11 (0.2-3 μm) and ASV555 (0.2-3 μm)). The color on top of the graphs indicates the occurrence category of each example (broad, intermediate and narrow based on their presence on the time series). B). Summary of the Wavelet Power Average of all ASVs that exhibited a significant recurrency in the wavelets analysis and the category they belong to based on both, the Wavelets Analyses and their occurrence over the time series (broad (>2/3), intermediate (1/3 – 2/3) and narrow (<1/3). Those plots highlighted inside the shadowed square belong to seasonal ASVs, first row, whereas the other three belong to chaotic ASVs based on the wavelets analysis, second row.

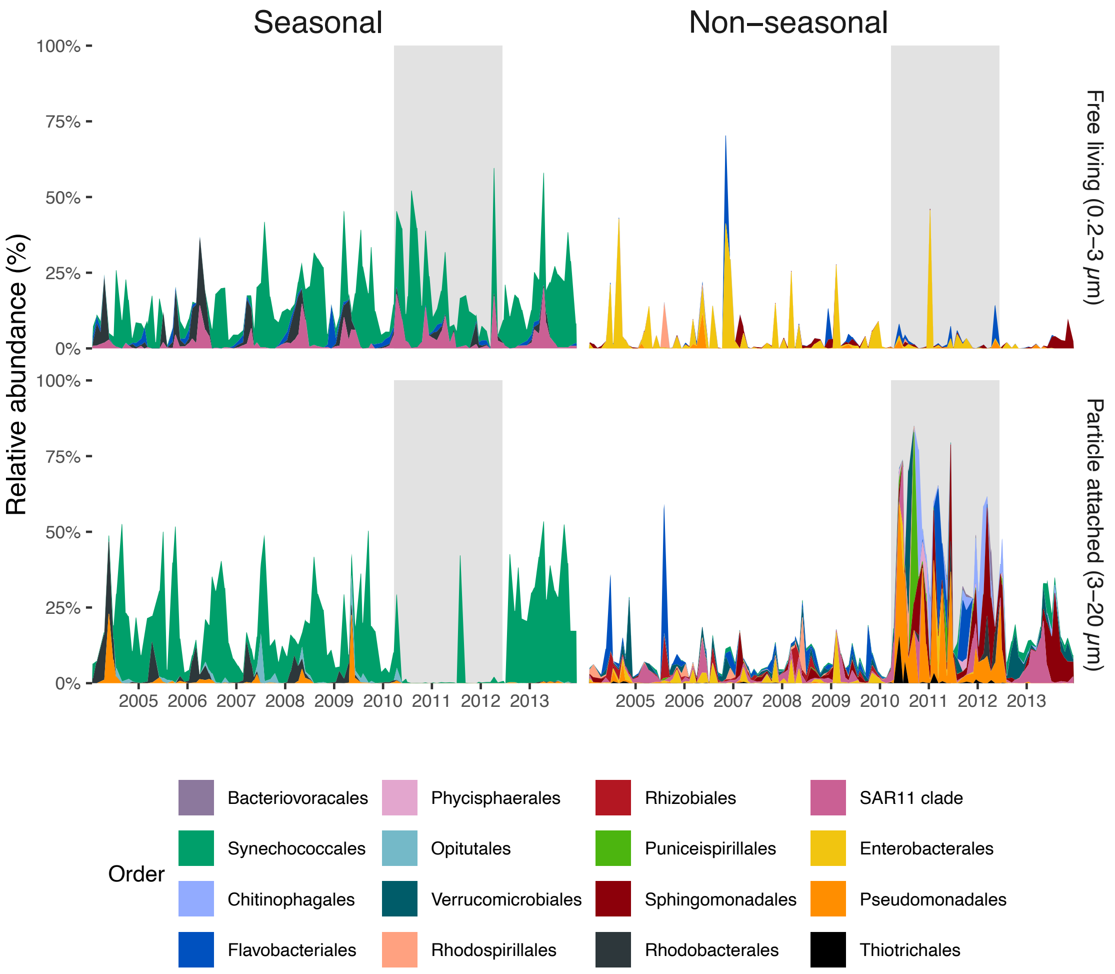

**Figure S7**: Blooming community in our time series in each size fraction (0.2-3 μm or 3-20 μm) and whether they presented seasonal or chaotic fluctuations over the years. The X-axis presents time and the Y-axis the relative abundance (%). The color denotes the taxonomic order each ASV belongs to, and the shaded area indicates the harbor restoration period.

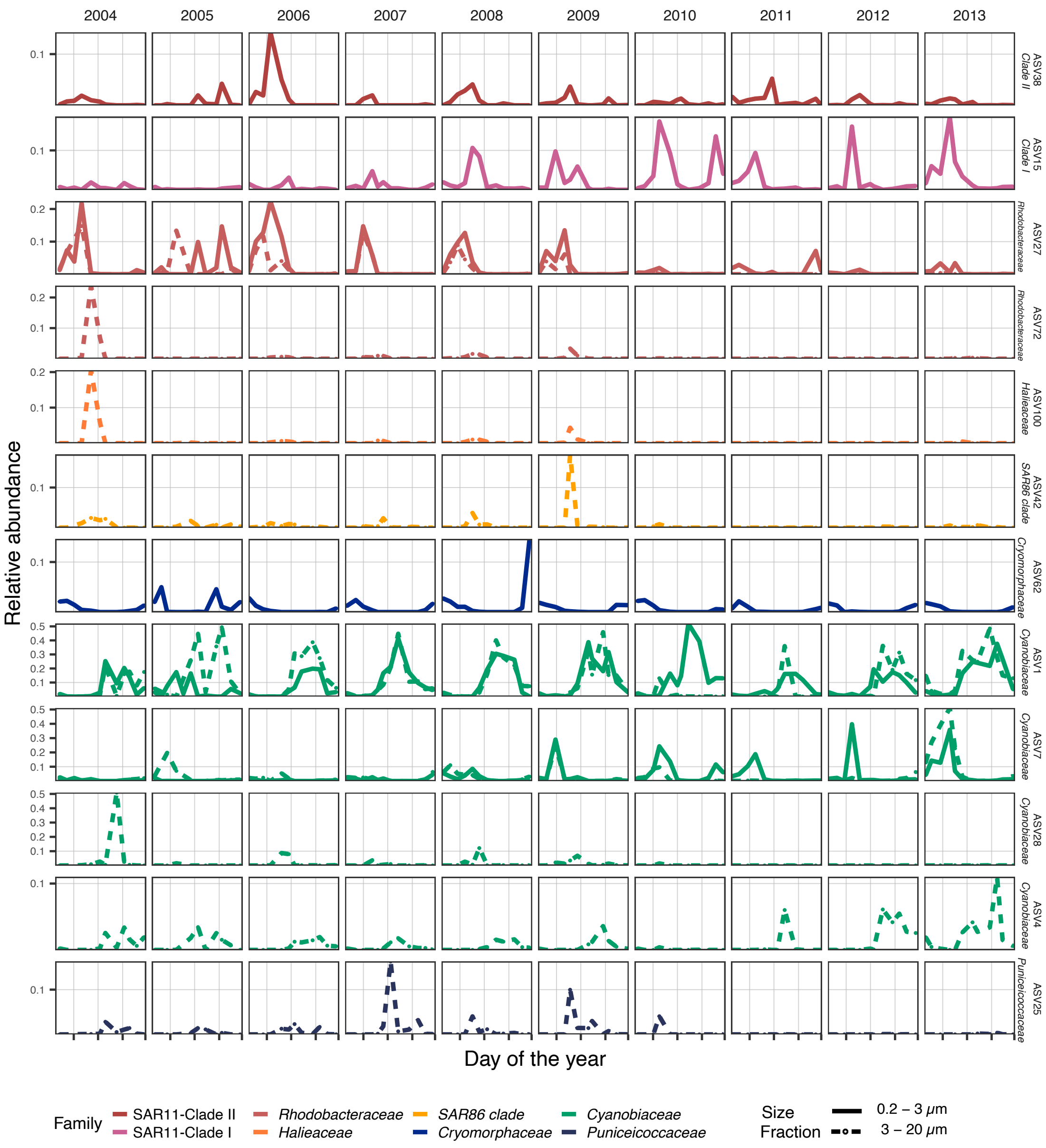

**Figure S8:** Seasonal bloomers and their interannual variability. The figure shows the different seasonal bloomers (determined with the wavelets analysis). The days of the year are in the X-axis, and the relative abundance of these taxa on the Y-axis. Each column represents a different year, and each row a different ASV. Note that Y-axes scales are different for each taxon. The line at 0.1 indicates the threshold used to determine a potential blooming event based on relative abundances, and the vertical lines indicate the different astronomical seasons of the year. The color indicates the taxonomic family of each ASV.

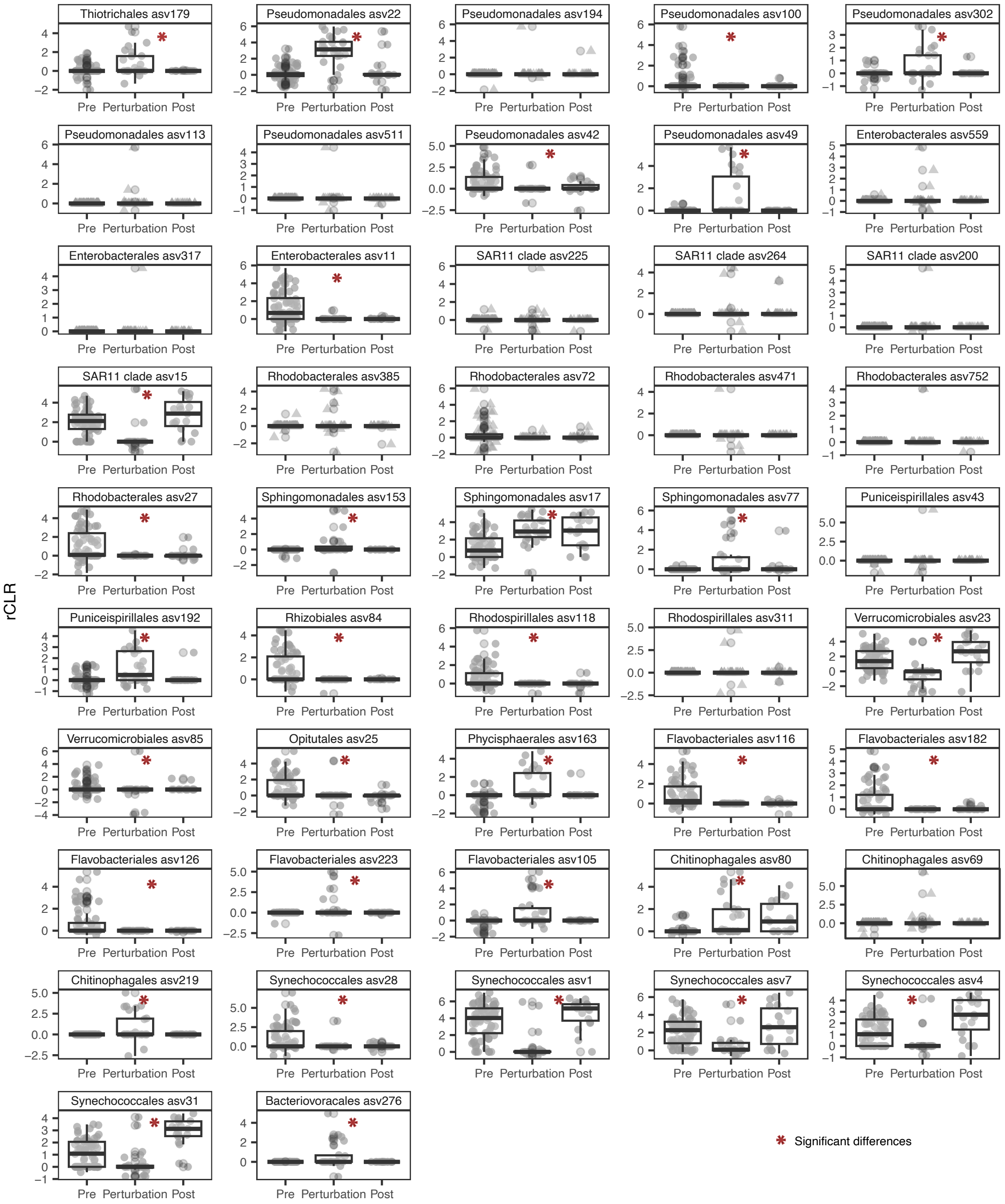

**Figure S9:** Statistical differences before, during and after the harbor restoration of bloomer ASVs abundances in the 3-20 µm size fraction (n = 47). Difference in bloomer abundances before (n = 72), during (n = 26), and after (n = 19) the identified perturbation (harbor restoration) in the 3-20 µm size fraction. Statistical differences between groups were assessed with ANOVA tests for normal data (Shapiro-Wilk test, $p > 0.01$) or were assessed using the Kruskal-Wallis test if non-normal (Shapiro-Wilk test, $p < 0.01$). Those differences during the harbor restoration that are significant are indicated with an asterisk.

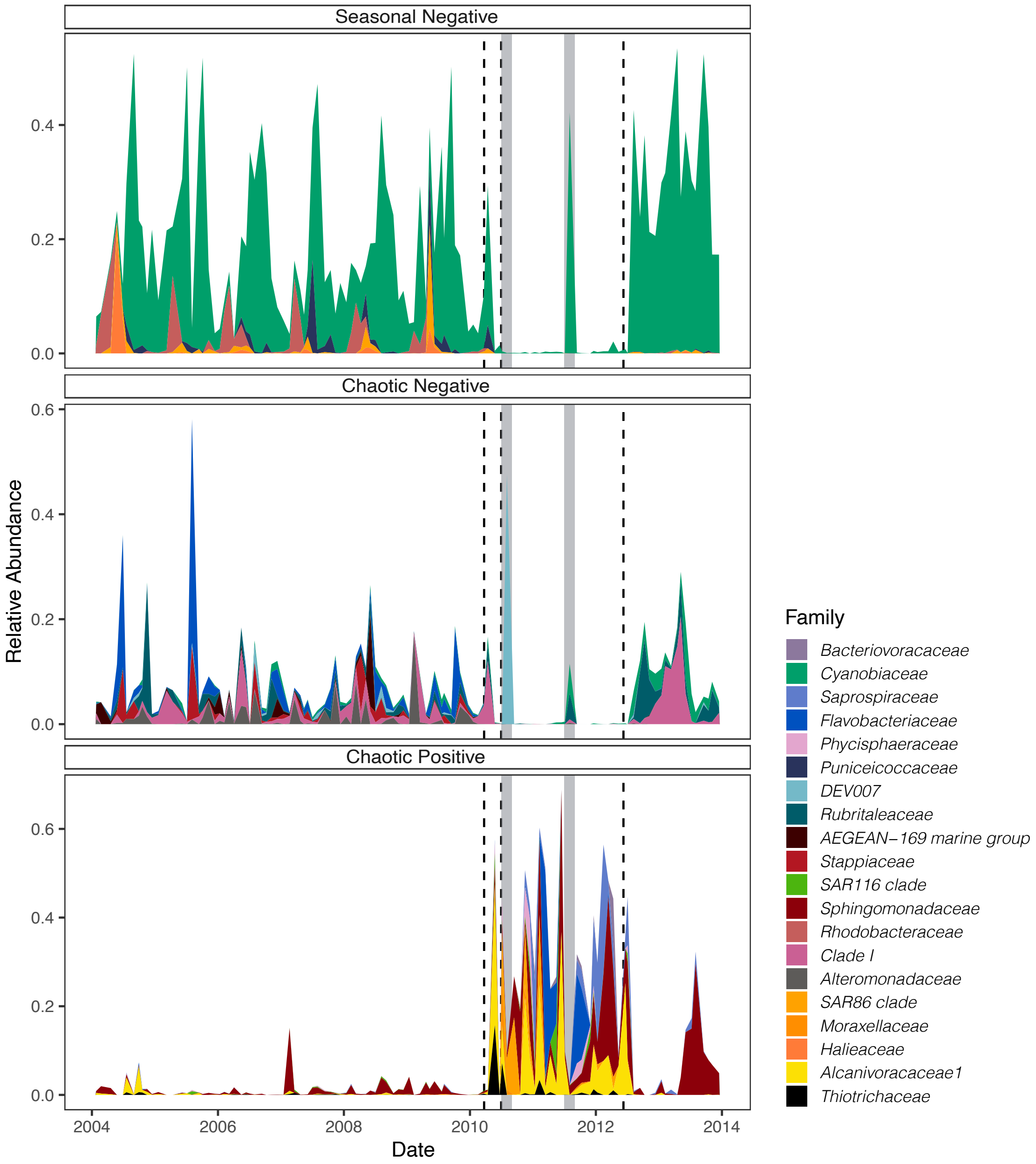

**Figure S10:** Relative abundance (%) of the different groups of ASVs affected by the harbor restoration in the 3-20 μm size fraction. Dashed lines represent the beginning of the harbor restoration period, the end of the most important works, and the complete end of the restoration. The shaded area in the time series correspond to the summer period during the harbor restoration, when the works stopped due to the high tourism in the area. Upper panel: seasonal blooming ASVs negatively affected by the perturbation, central panel: chaotic blooming ASVs negatively affected by the perturbation, and lower panel: chaotic blooming ASVs positively affected by the perturbation.

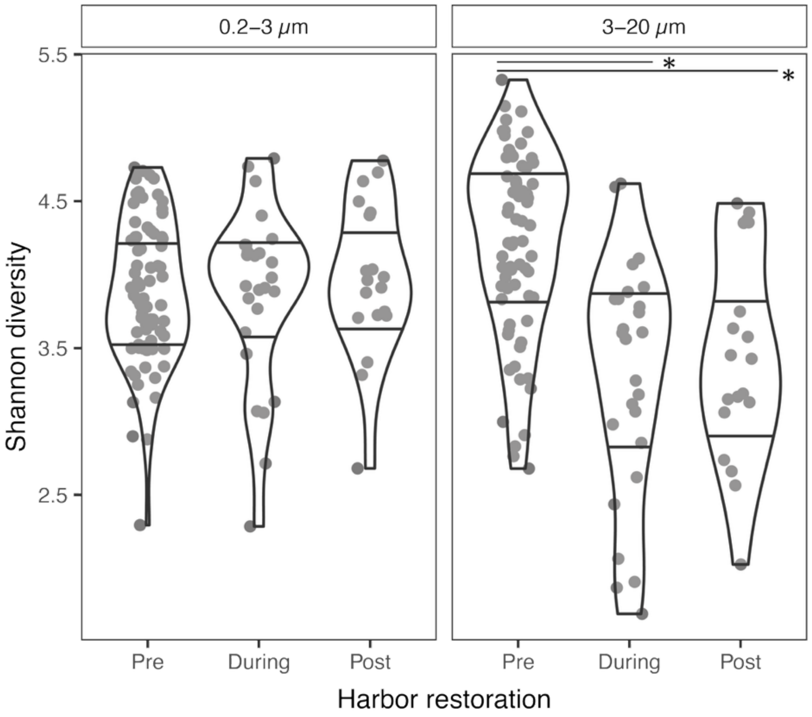

**Figure S11:** Bacterial diversity changes during harbor restoration. Shannon diversity in the samples before (n = 75 (0.2–3 μm), n = 72 (3-20 μm)), during (n = 26, for each fraction), and after (n = 19, for each fraction), the restoration of the Blanes harbor. Statistical differences between groups were assessed using the Kruskal-Wallis test due to non-normality (Shapiro-Wilk test, $p < 0.01$).

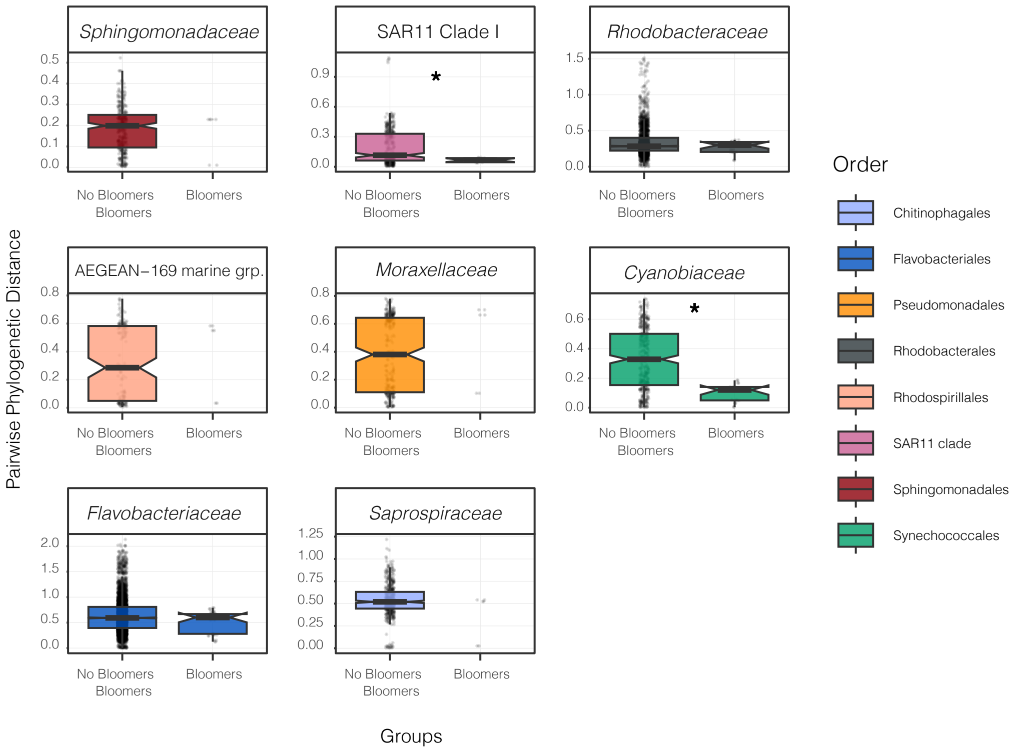

**Figure S12:** Boxplots comparing pairwise phylogenetic distances between blooming and non-blooming ASVs, and among blooming ASVs, across taxonomic families containing more than one blooming taxon. Groups without boxplots reflect insufficient numbers of pairwise comparisons. Significant differences were assessed using a non-parametric Wilcoxon test and are highlighted with an asterisk. The color of the boxplots indicates the phylogenetic Order.

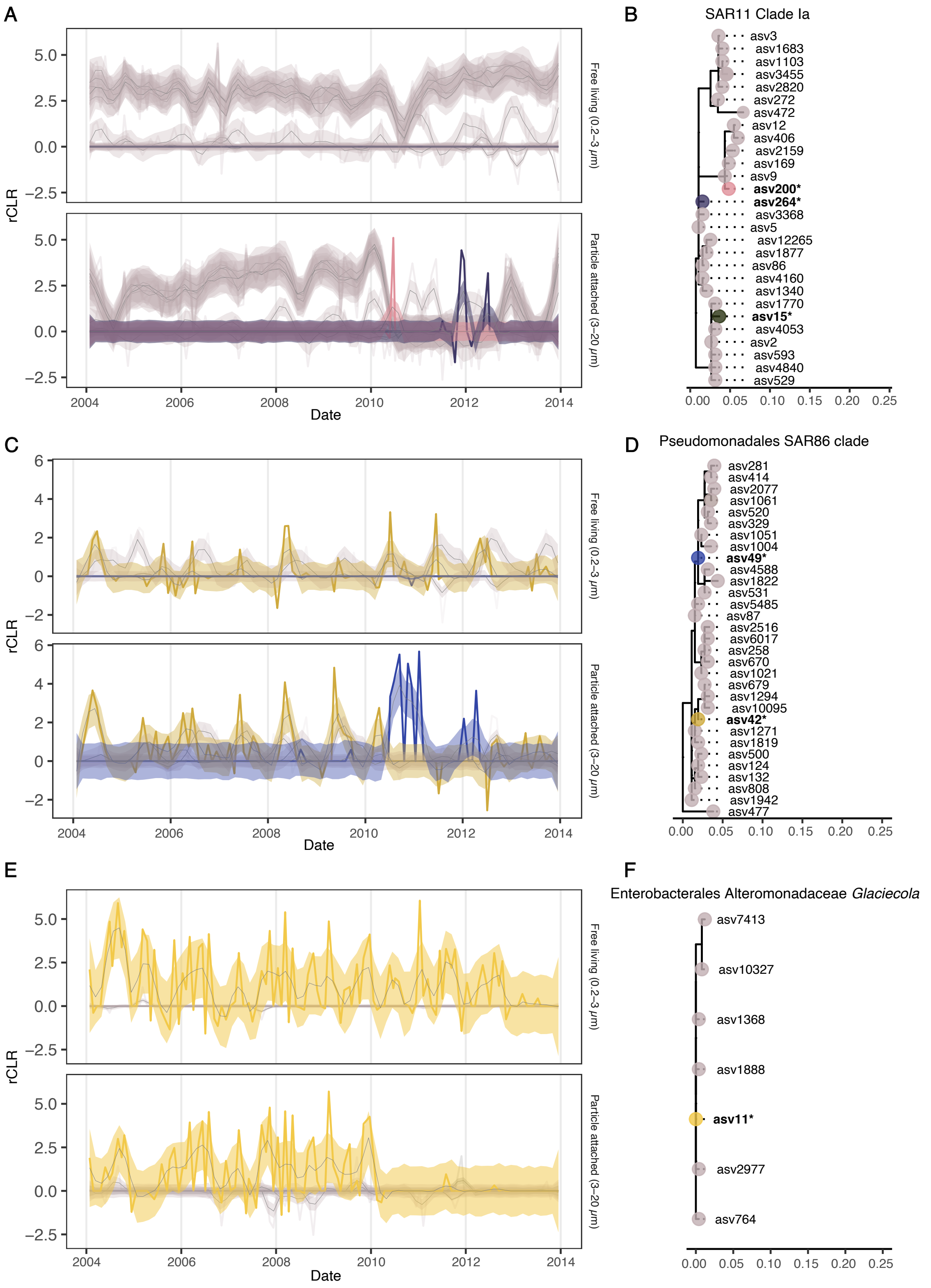

**Figure S13:** Examples of the time-series of closely related bloomers (in bright colors) and their closely related ASVs (p-distance < 0.012, in dimmed colors) in the SAR11 clade (A), SAR86 Clade (C) and genus *Glaciecola* (E). Phylogenetic trees of the closely related ASVs in the SAR11 clade (B), SAR86 Clade (D) and genus *Glaciecola* (F). Those ASVs that exhibited a bloomer behavior in our time series are highlighted in bold and with an asterisk.

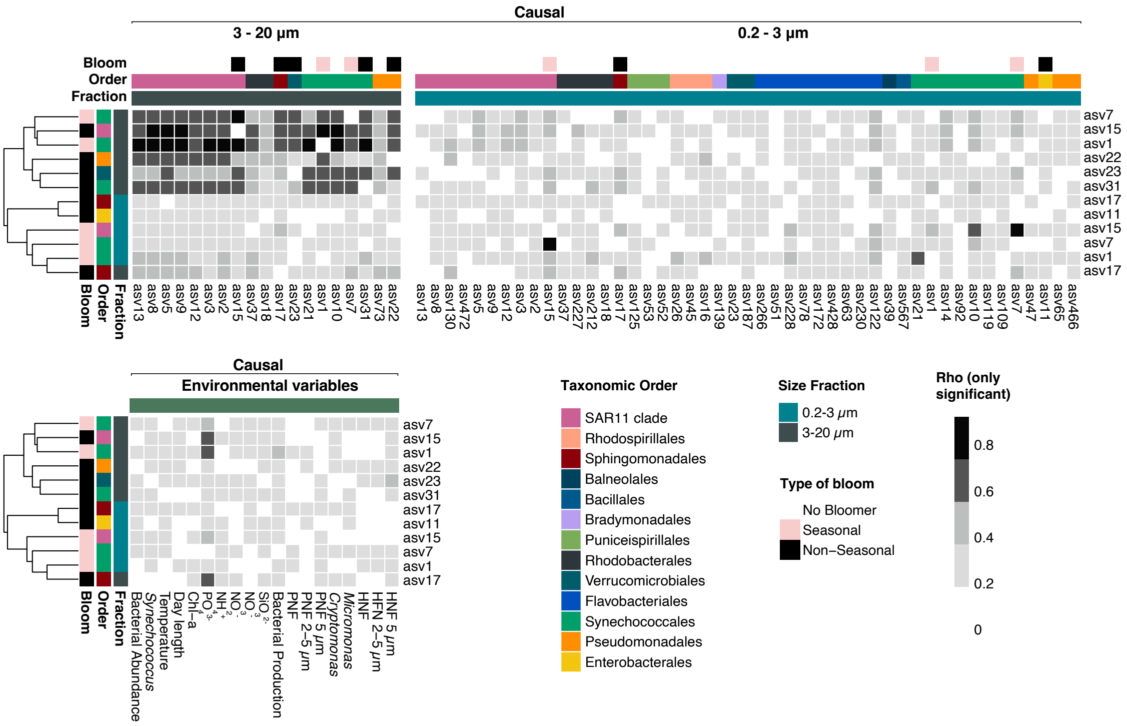

**Figure S14:** Convergent Cross Mapping (CCM) results showing the causality of different ASVs and environmental variables on the bloomers. Only significant values are presented. The columns represent causal variables affecting the rows (i.e., the bloomers), with higher rho values indicating stronger predictive skill, a measure of strength of causality on the bloomers' abundances. The environmental variables included were selected for having minimal missing data: PNF: total phototrophic nanoflagellates, PNF 2-5 µm: phototrophic nanoflagellates (2-5 µm), PNF 5 µm: phototrophic nanoflagellates (>5 µm); HNF: total heterotrophic nanoflagellates, HNF 2-5 µm: heterotrophic nanoflagellates (2-5 µm), HNF 5 µm: heterotrophic nanoflagellates (>5 µm); Total bacterial abundance (cells/mL), *Synechococcus abundance* (cells/mL); temperature (°C), day length (hours), Chl-a: chlorophyll *a* (µg/L); nutrient concentrations (µM): phosphate ($PO_4^{3-}$), ammonia ($NH_4^+$), nitrite ($NO_2^-$), nitrate ($NO_3^-$), and silicate ($SiO_3^{2-}$); bacterial production (µg $C \cdot L^{-1} \cdot d^{-1}$); *Cryptomonas* (cells/mL), and *Micromonas-like phototrophs* (cells/mL).

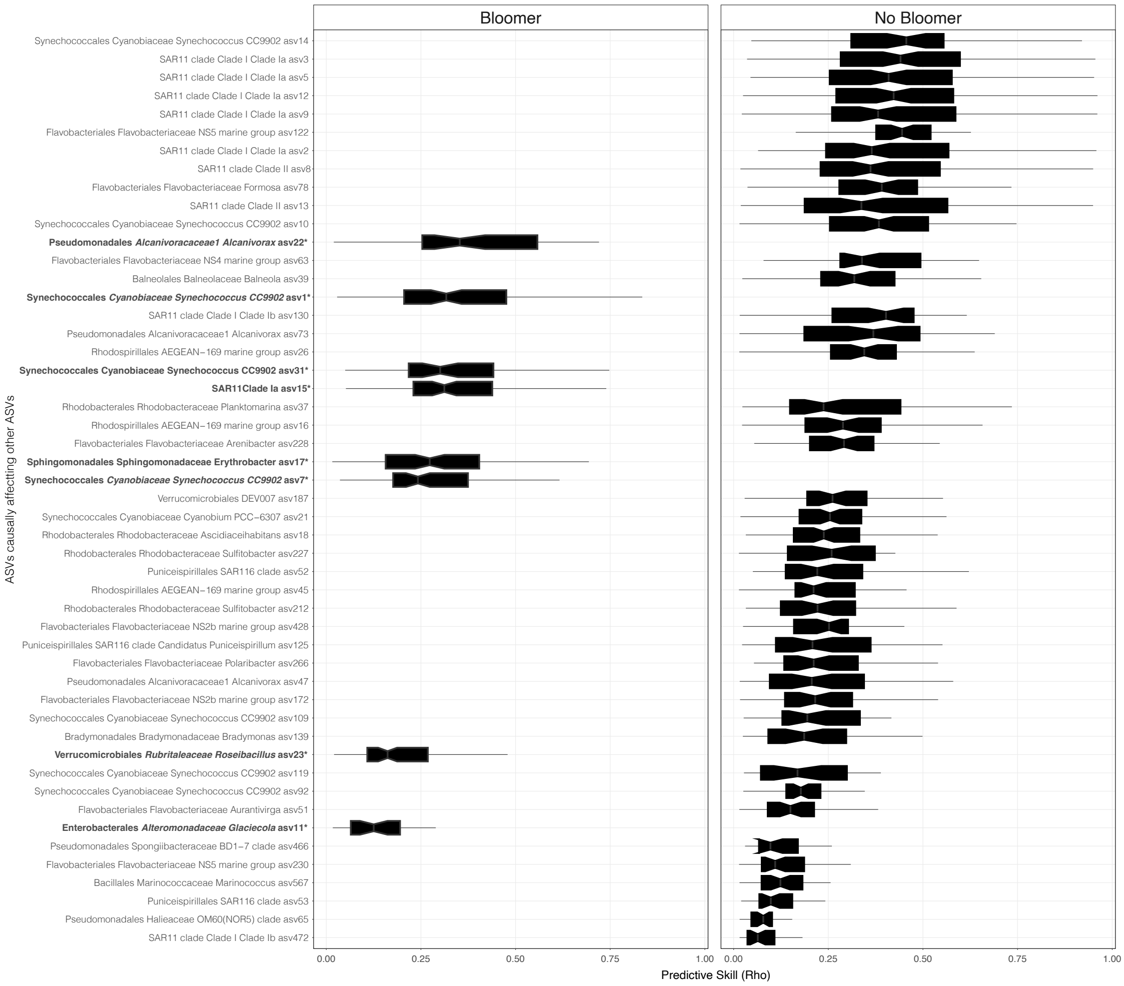

**Figure S15:** Boxplot of occurrent ASVs (> 1/3 occurrence) ordered by their causal effect on other ASVs in the community, identified using Convergent Cross Mapping (CCM). ASVs are ordered by the increasing number of significative predictive skills (rho) values obtained. The left panel corresponds to those ASVs identified as bloomers in our time series, with their taxonomy in bold and labelled with an asterisk in the Y axis.

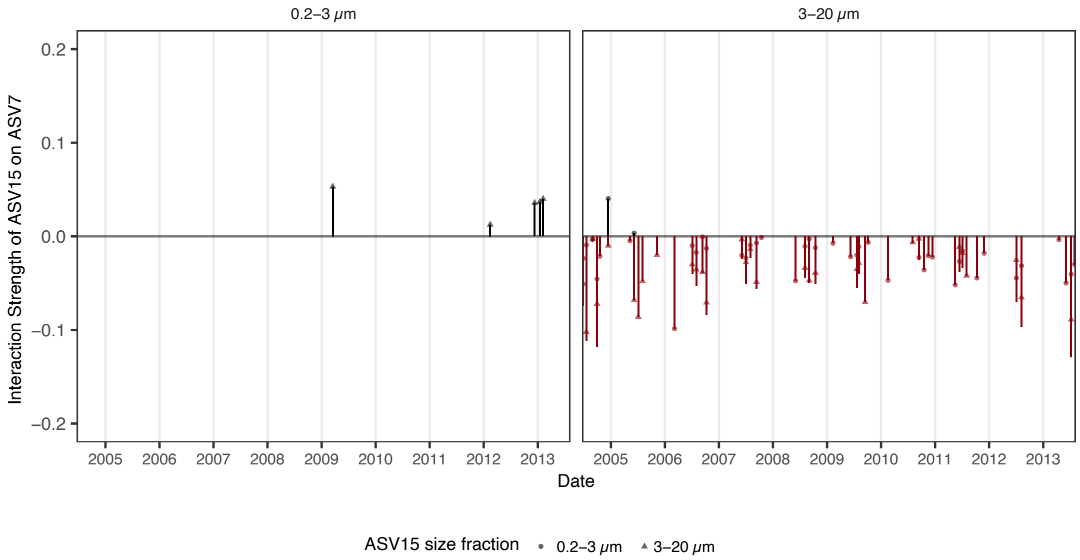

**Figure S16:** Multiview distance regularized (MDR) S-map presenting the interaction strength that SAR 11 Clade Ia–ASV15 has on *Synechococcus*–ASV7 during the time series. The symbol shape indicates the fraction in which ASV15 is affecting ASV7 in either the 0.2 – 3 µm (free-living) size fraction or the 3-20 µm (particle-attached) one.
