## Supplementary Information II for "The natural history of bacterial bloomers in a decade-long time series"

### Index

### Wavelets spectra supplementary figures

#### Shared ASVs between 3-20 µm and 0.2-3 µm size fractions

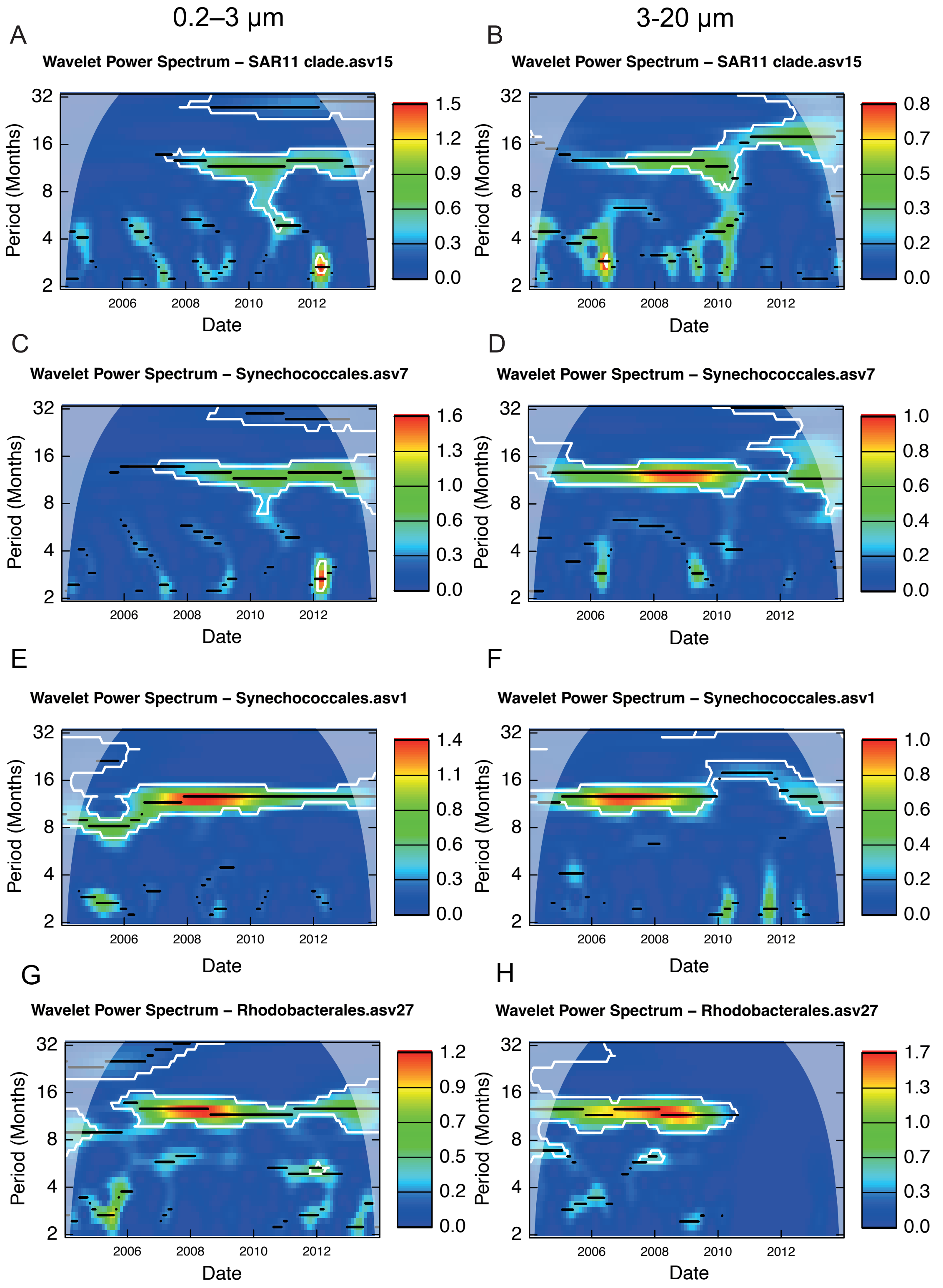

**Figure S2.1.** Continuous wavelets transformation for ASV present in both size fractions. Each plot corresponds to a different taxon. The X-axis represents the time series (2004-2014), and the Y-axis shows the period (in months). Color intensity indicates wavelet power, reflecting the strength of periodic signals. Significant (p-value after consistent results with 100 repetitions) signals within a period are highlighted by contour lines, while the color scale suggests that a periodic phenomenon is detected in the pattern of abundance of each ASV at that period. Stronger signal intensities at specific periods suggest recurrent patterns in ASV abundance. Temporal scales are interpreted as: short-term (2–4 months), semiannual (4–8 months), seasonal (8–16 months), annual (16–32 months), and interannual (~32 months). In case that an ASV has a recurrent pattern in one period we would observe a higher signal intensity at that period. The plots were generated with the package *WaveletComp*, which uses a Morlet wavelet.

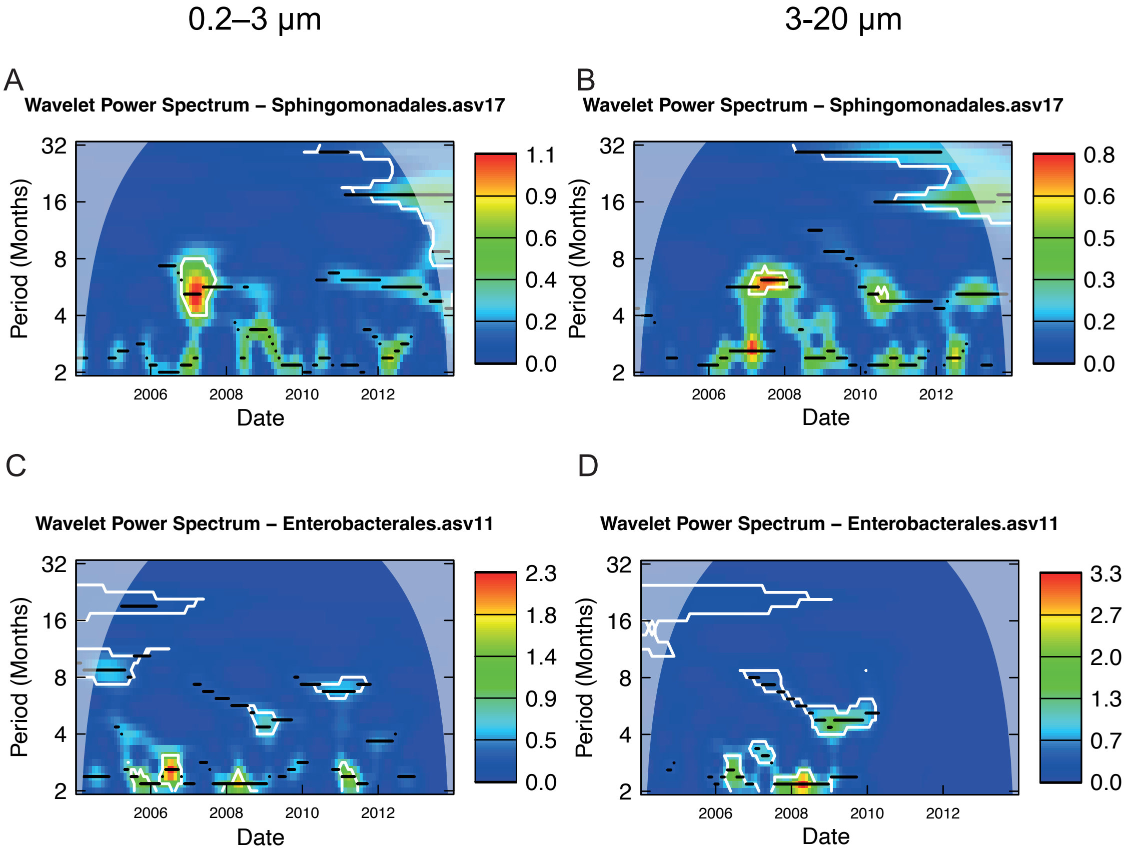

**Figure S2.2.** Continuous wavelets transformation for ASV present in both size fractions. Each plot corresponds to a different taxon. The X-axis represents the time series (2004-2014), and the Y-axis shows the period (in months). Color intensity indicates wavelet power, reflecting the strength of periodic signals. Significant (p-value after consistent results with 100 repetitions) signals within a period are highlighted by contour lines, while the color scale suggests that a periodic phenomenon is detected in the pattern of abundance of each ASV at that period. Stronger signal intensities at specific periods suggest recurrent patterns in ASV abundance. Temporal scales are interpreted as: short-term (2–4 months), semiannual (4–8 months), seasonal (8–16 months), annual (16–32 months), and interannual (~32 months). In case that an ASV has a recurrent pattern in one period we would observe a higher signal intensity at that period. The plots were generated with the package *WaveletComp*, which uses a Morlet wavelet.

## 0.2–3 µm

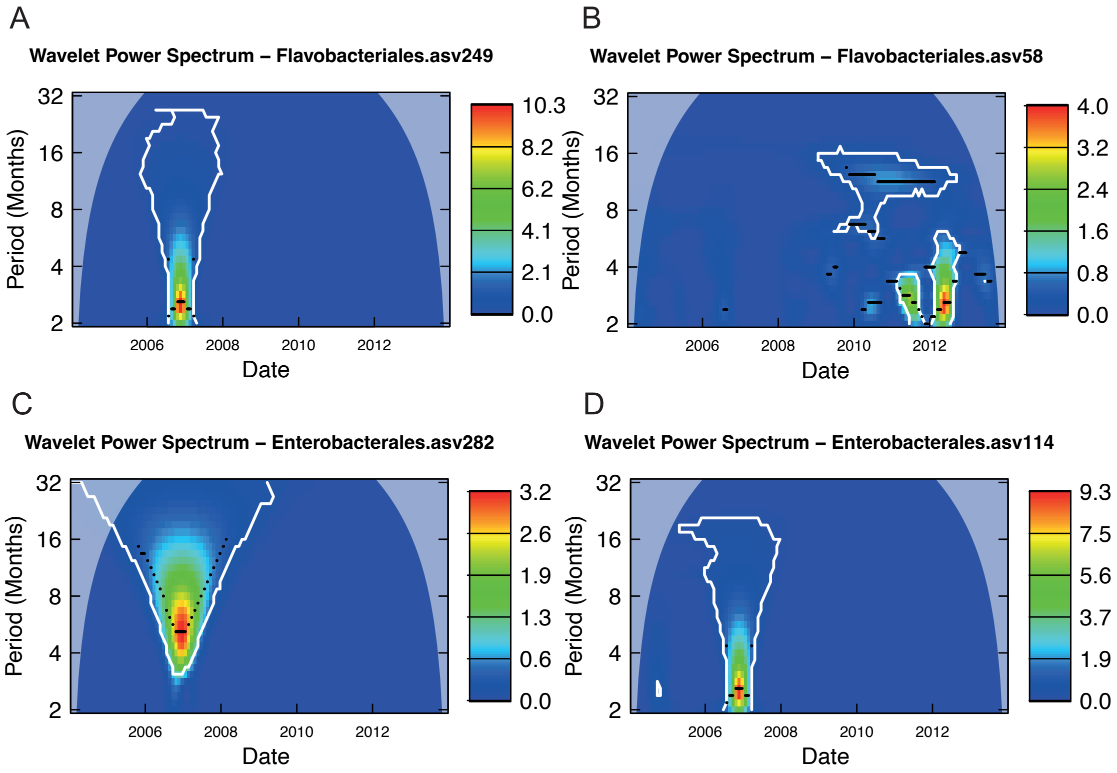

**Figure S2.3.** Continuous wavelets transformation for ASV present in the free-living fraction (0.2-3 µm). Each plot corresponds to a different taxon. The X-axis represents the time series (2004-2014), and the Y-axis shows the period (in months). Color intensity indicates wavelet power, reflecting the strength of periodic signals. Significant (p-value after consistent results with 100 repetitions) signals within a period are highlighted by contour lines, while the color scale suggests that a periodic phenomenon is detected in the pattern of abundance of each ASV at that period. Stronger signal intensities at specific periods suggest recurrent patterns in ASV abundance. Temporal scales are interpreted as: short-term (2–4 months), semiannual (4–8 months), seasonal (8–16 months), annual (16–32 months), and interannual (~32 months). In case that an ASV has a recurrent pattern in one period we would observe a higher signal intensity at that period. The plots were generated with the package *WaveletComp*, which uses a Morlet wavelet.

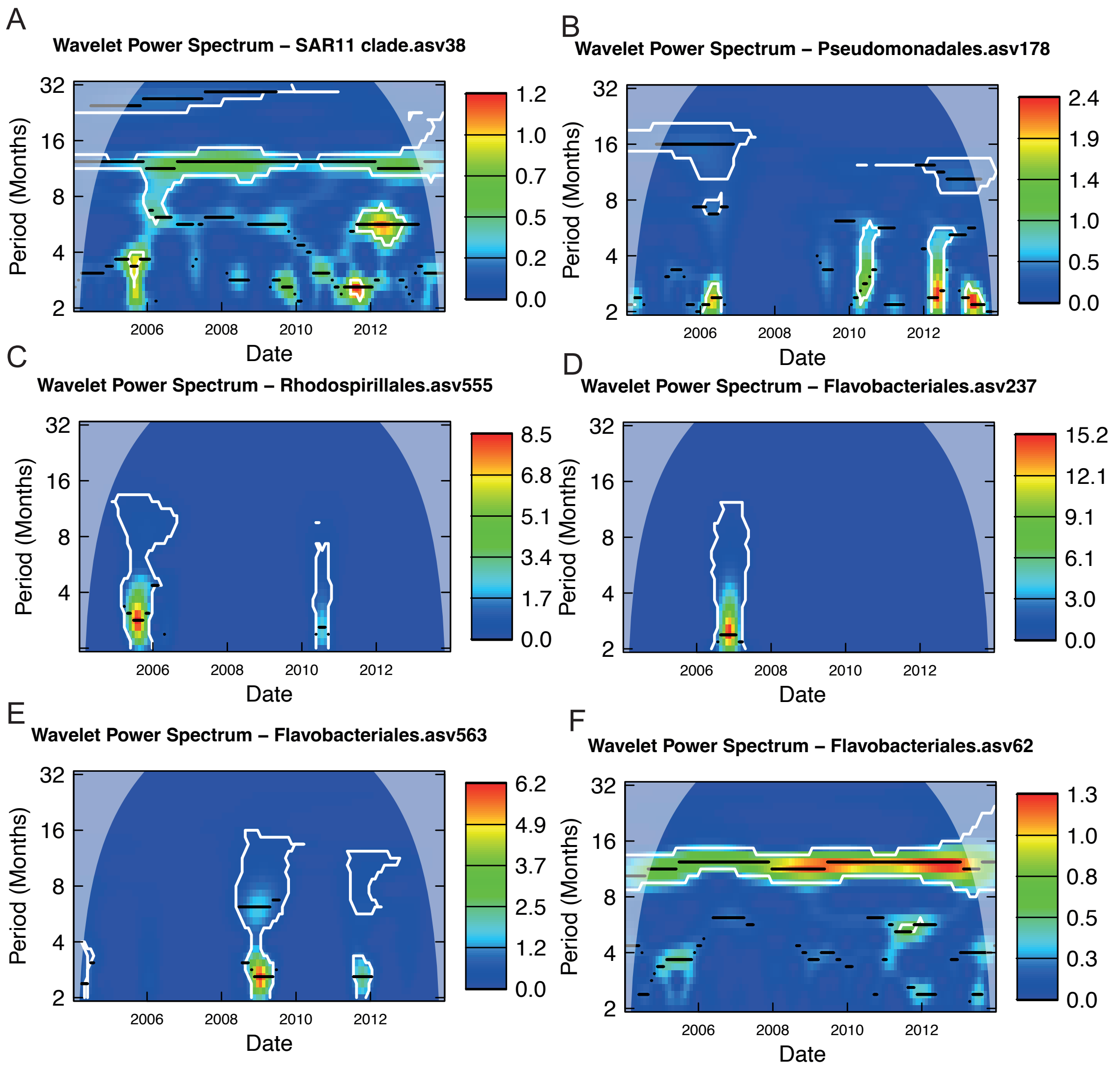

**Figure S2.4.** Continuous wavelets transformation for ASV present in the free-living fraction (0.2-3 μm). Each plot corresponds to a different taxon. The X-axis represents the time series (2004-2014), and the Y-axis shows the period (in months). Color intensity indicates wavelet power, reflecting the strength of periodic signals. Significant (p-value after consistent results with 100 repetitions) signals within a period are highlighted by contour lines, while the color scale suggests that a periodic phenomenon is detected in the pattern of abundance of each ASV at that period. Stronger signal intensities at specific periods suggest recurrent patterns in ASV abundance. Temporal scales are interpreted as: short-term (2–4 months), semiannual (4–8 months), seasonal (8–16 months), annual (16–32 months), and interannual (~32 months). In case that an ASV has a recurrent pattern in one period we would observe a higher signal intensity at that period. The plots were generated with the package *WaveletComp*, which uses a Morlet wavelet.

## 3–20 µm

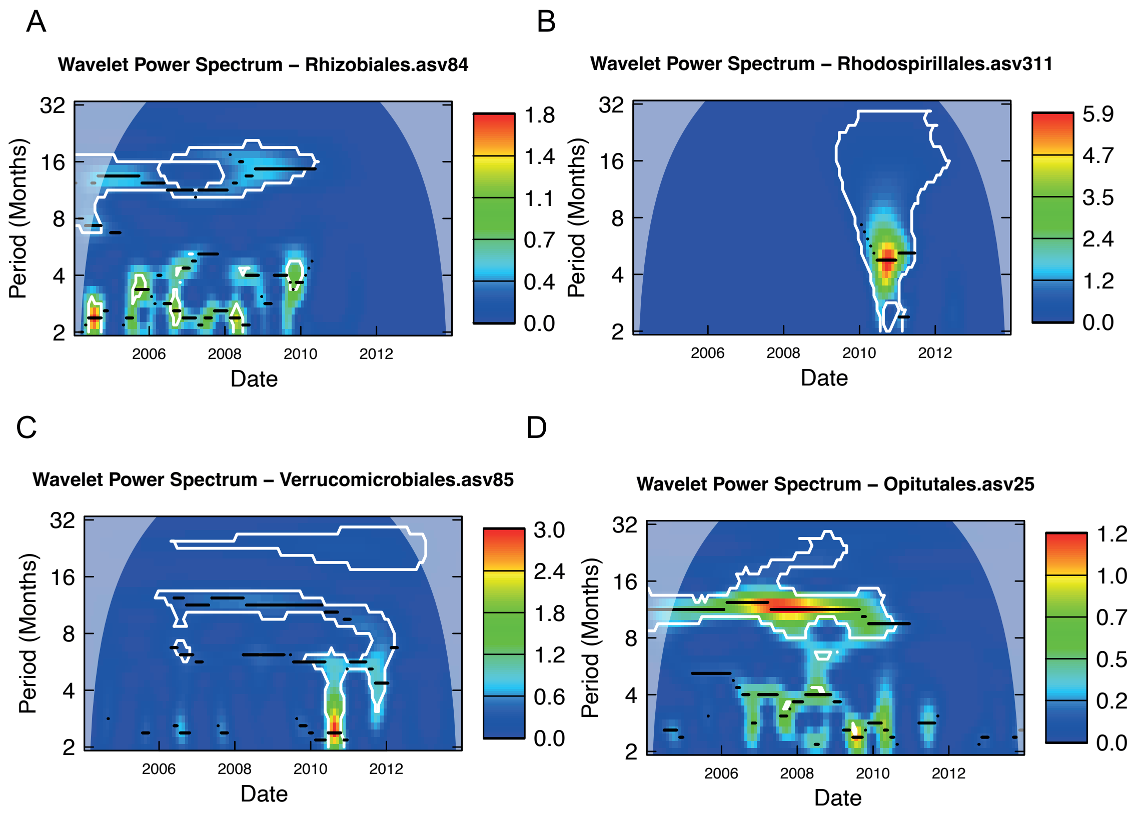

**Figure S2.5.** Continuous wavelets transformation for ASV present in the Particle-attached fraction (3-20 µm) size fraction. Each plot corresponds to a different taxon. The X-axis represents the time series (2004-2014), and the Y-axis shows the period (in months). Color intensity indicates wavelet power, reflecting the strength of periodic signals. Significant (p-value after consistent results with 100 repetitions) signals within a period are highlighted by contour lines, while the color scale suggests that a periodic phenomenon is detected in the pattern of abundance of each ASV at that period. Stronger signal intensities at specific periods suggest recurrent patterns in ASV abundance. Temporal scales are interpreted as: short-term (2–4 months), semiannual (4–8 months), seasonal (8–16 months), annual (16–32 months), and interannual (~32 months). In case that an ASV has a recurrent pattern in one period we would observe a higher signal intensity at that period. The plots were generated with the package *WaveletComp*, which uses a Morlet wavelet.

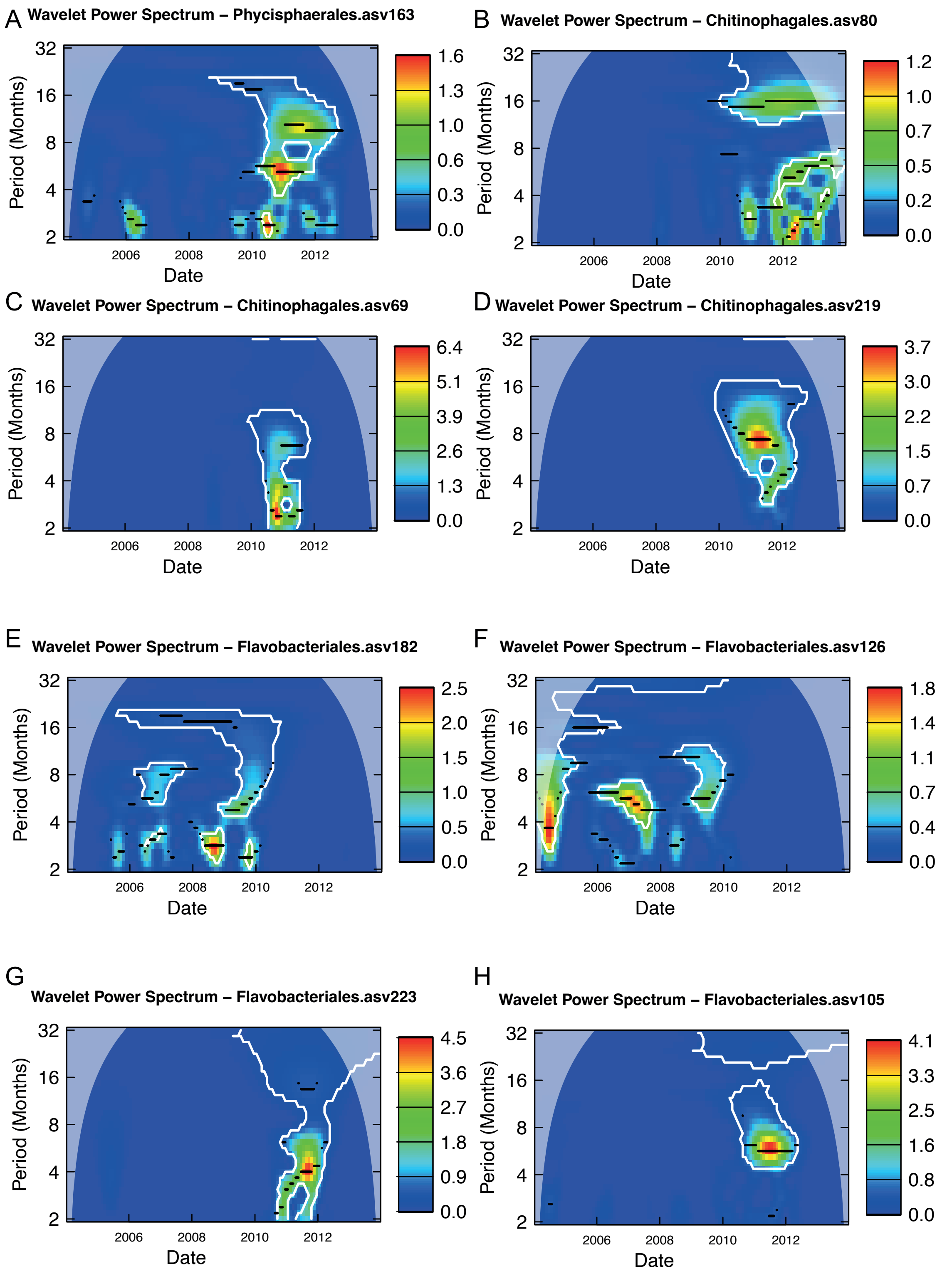

**Figure S2.6.** Continuous wavelets transformation for ASV present in the Particle-attached fraction (3-20 μm). Each plot corresponds to a different taxon. The X-axis represents the time series (2004-2014), and the Y-axis shows the period (in months). Color intensity indicates wavelet power, reflecting the strength of periodic signals. Significant (p-value after consistent results with 100 repetitions) signals within a period are highlighted by contour lines, while the color scale suggests that a periodic phenomenon is detected in the pattern of abundance of each ASV at that period. Stronger signal intensities at specific periods suggest recurrent patterns in ASV abundance. Temporal scales are interpreted as: short-term (2–4 months), semiannual (4–8 months), seasonal (8–16 months), annual (16–32 months), and interannual (~32 months). In case that an ASV has a recurrent pattern in one period we would observe a higher signal intensity at that period. The plots were generated with the package *WaveletComp*, which uses a Morlet wavelet.

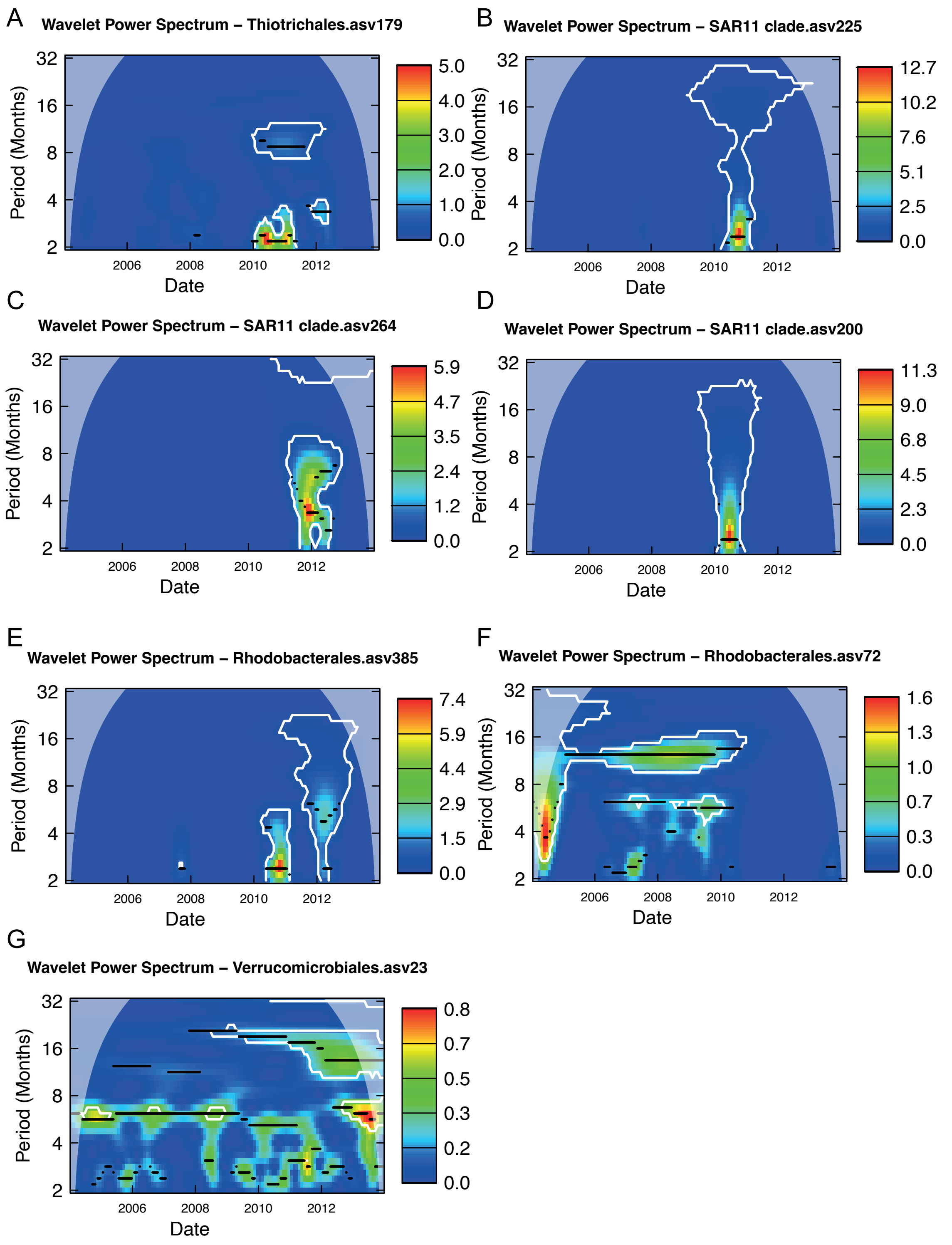

**Figure S2.7.** Continuous wavelets for ASV present in the Particle-attached fraction (3-20 μm Each plot corresponds to a different taxon. The X-axis represents the time series (2004-2014), and the Y-axis shows the period (in months). Color intensity indicates wavelet power, reflecting the strength of periodic signals. Significant (p-value after consistent results with 100 repetitions) signals within a period are highlighted by contour lines, while the color scale suggests that a periodic phenomenon is detected in the pattern of abundance of each ASV at that period. Stronger signal intensities at specific periods suggest recurrent patterns in ASV abundance. Temporal scales are interpreted as: short-term (2–4 months), semiannual (4–8 months), seasonal (8–16 months), annual (16–32 months), and interannual (~32 months). In case that an ASV has a recurrent pattern in one period we would observe a higher signal intensity at that period. The plots were generated with the package *WaveletComp*, which uses a Morlet wavelet.

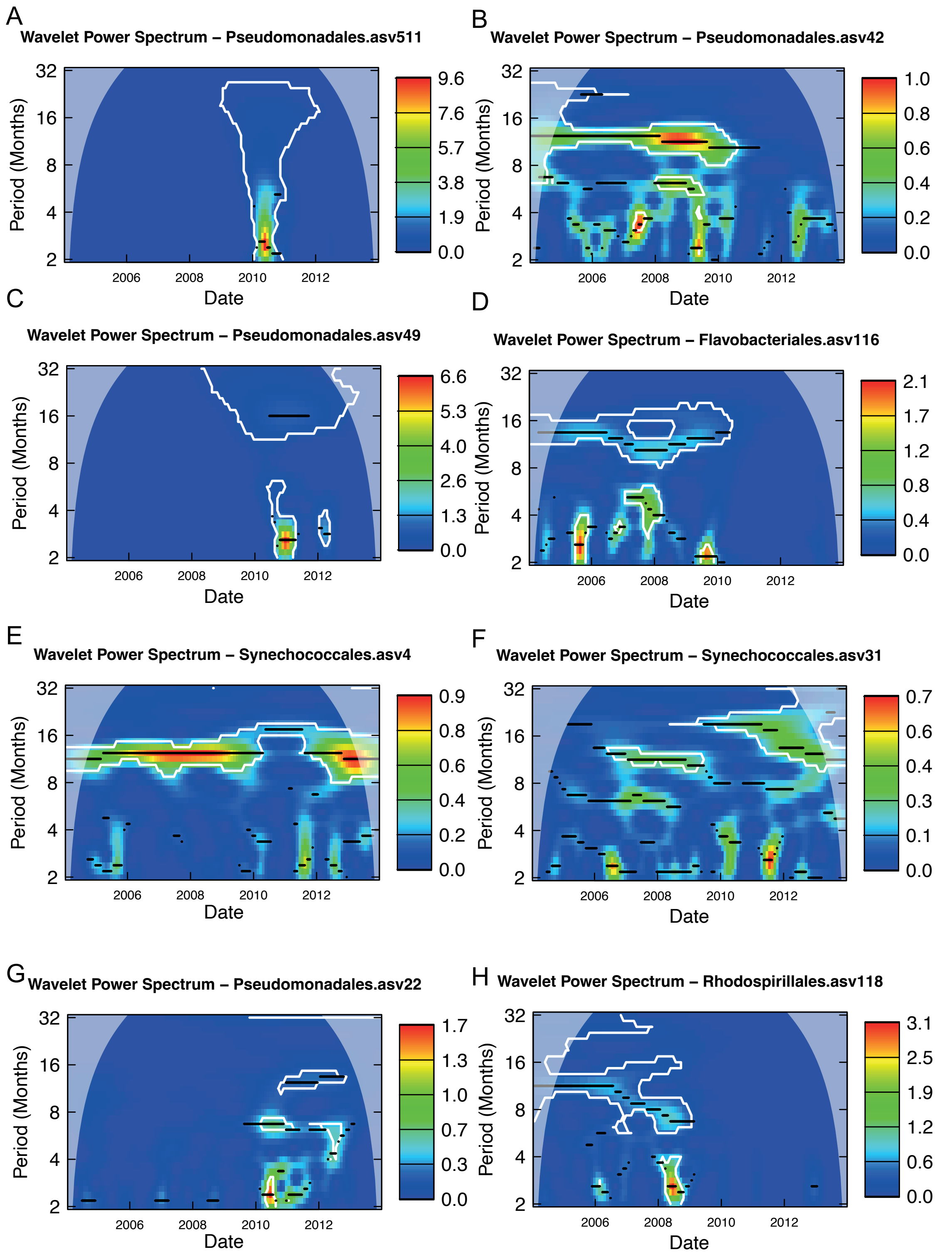

**Figure S2.8.** Continuous wavelets transformation for ASV present in the Particle-attached fraction (3-20 μm). Each plot corresponds to a different taxon. The X-axis represents the time series (2004-2014), and the Y-axis shows the period (in months). Color intensity indicates wavelet power, reflecting the strength of periodic signals. Significant (p-value after consistent results with 100 repetitions) signals within a period are highlighted by contour lines, while the color scale suggests that a periodic phenomenon is detected in the pattern of abundance of each ASV at that period. Stronger signal intensities at specific periods suggest recurrent patterns in ASV abundance. Temporal scales are interpreted as: short-term (2–4 months), semiannual (4–8 months), seasonal (8–16 months), annual (16–32 months), and interannual (~32 months). In case that an ASV has a recurrent pattern in one period we would observe a higher signal intensity at that period. The plots were generated with the package *WaveletComp*, which uses a Morlet wavelet.

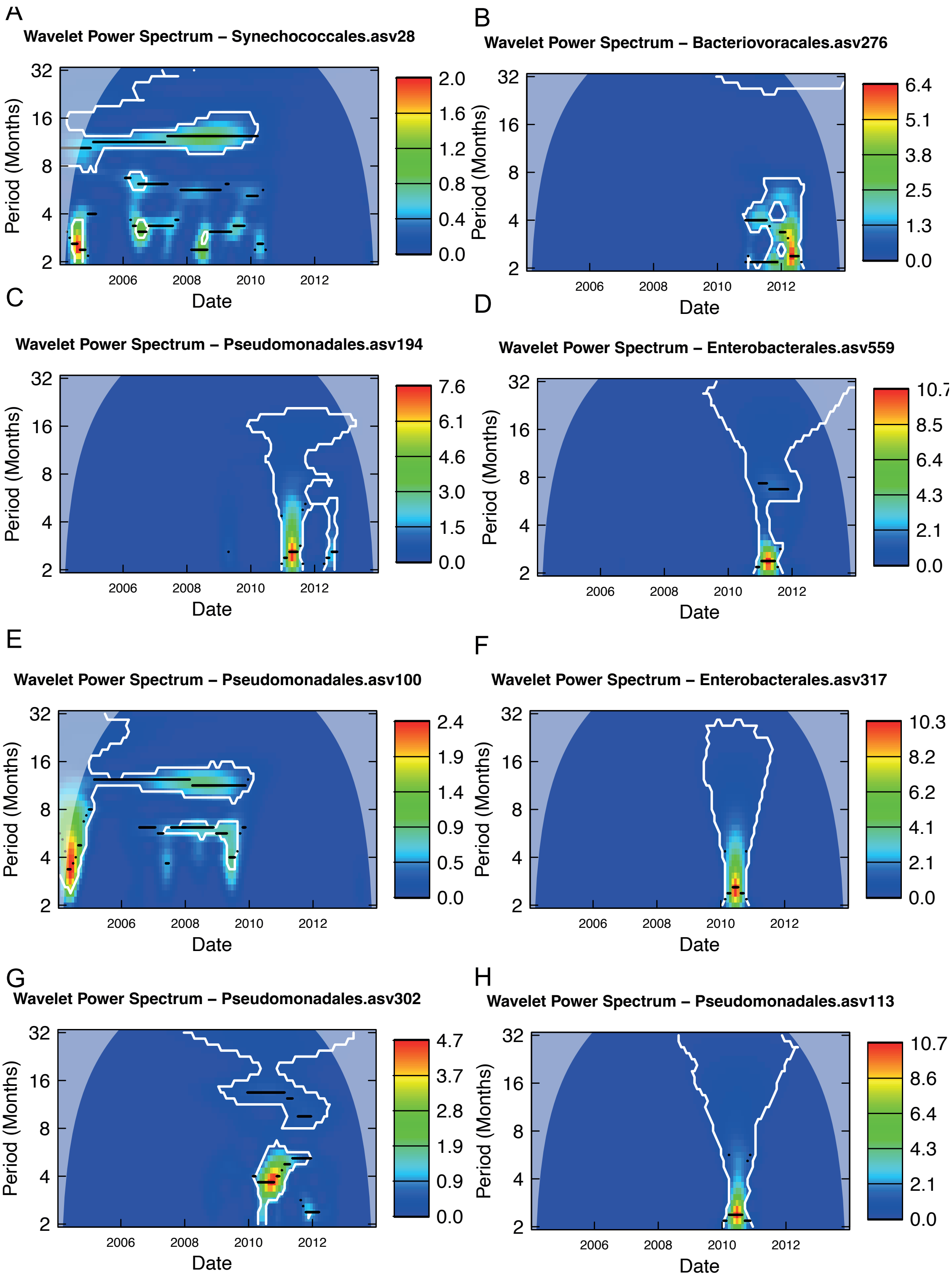

**Figure S2.9.** Continuous wavelets transformation for ASV present in the Particle-attached fraction (3-20 μm). Each plot corresponds to a different taxon. The X-axis represents the time series (2004-2014), and the Y-axis shows the period (in months). Color intensity indicates wavelet power, reflecting the strength of periodic signals. Significant (p-value after consistent results with 100 repetitions) signals within a period are highlighted by contour lines, while the color scale suggests that a periodic phenomenon is detected in the pattern of abundance of each ASV at that period. Stronger signal intensities at specific periods suggest recurrent patterns in ASV abundance. Temporal scales are interpreted as: short-term (2–4 months), semiannual (4–8 months), seasonal (8–16 months), annual (16–32 months), and interannual (~32 months). In case that an ASV has a recurrent pattern in one period we would observe a higher signal intensity at that period. The plots were generated with the package *WaveletComp*, which uses a Morlet wavelet.

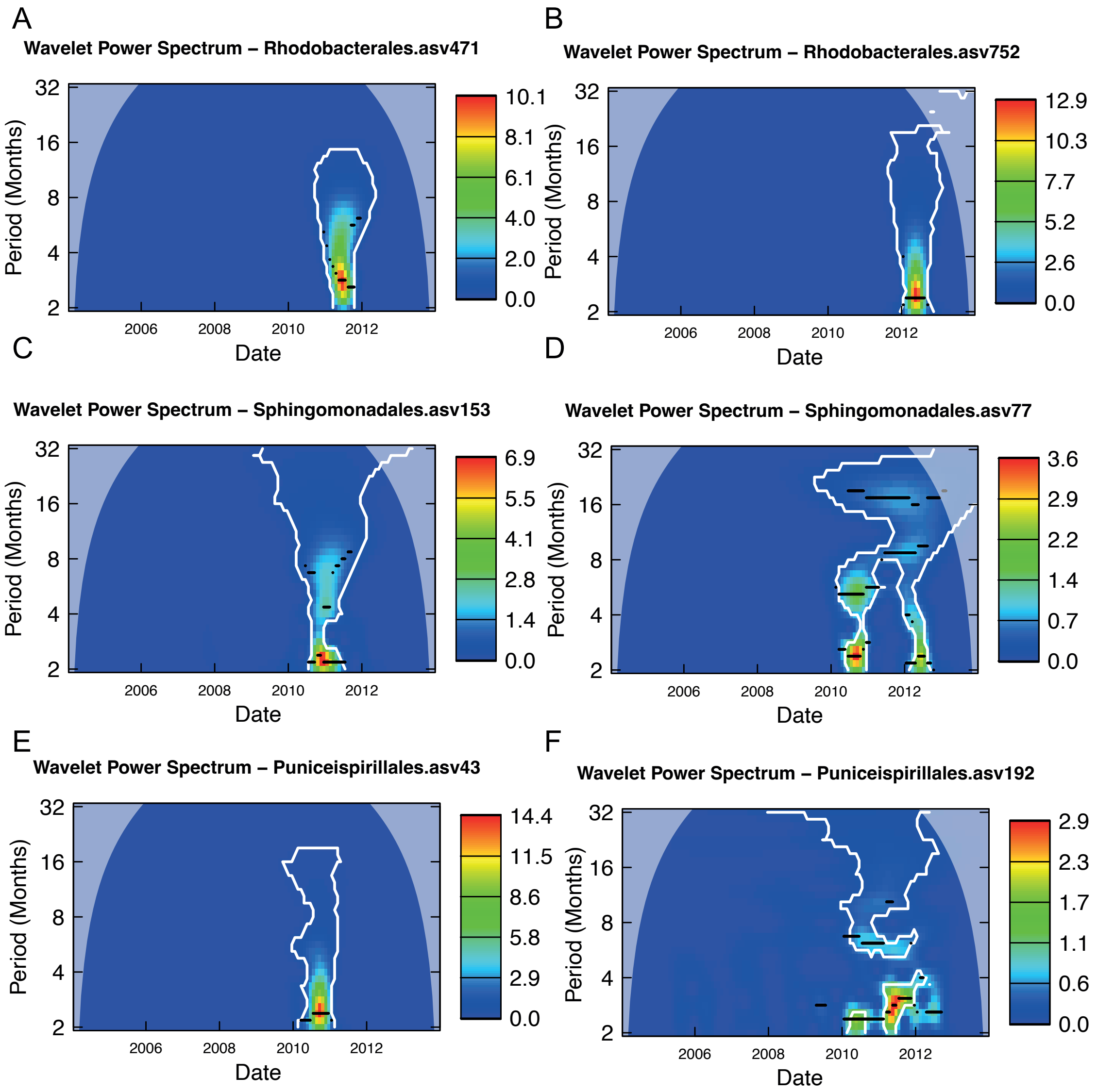

**Figure S2.10.** Continuous wavelets transformation for ASV present in the Particle-attached fraction (3-20 μm). Each plot corresponds to a different taxon. The X-axis represents the time series (2004-2014), and the Y-axis shows the period (in months). Color intensity indicates wavelet power, reflecting the strength of periodic signals. Significant (p-value after consistent results with 100 repetitions) signals within a period are highlighted by contour lines, while the color scale suggests that a periodic phenomenon is detected in the pattern of abundance of each ASV at that period. Stronger signal intensities at specific periods suggest recurrent patterns in ASV abundance. Temporal scales are interpreted as: short-term (2–4 months), semiannual (4–8 months), seasonal (8–16 months), annual (16–32 months), and interannual (~32 months). In case that an ASV has a recurrent pattern in one period we would observe a higher signal intensity at that period. The plots were generated with the package *WaveletComp*, which uses a Morlet wavelet.
